# Life-history trade-offs explain local adaptation in *Arabidopsis thaliana*

**DOI:** 10.1101/2025.05.18.654693

**Authors:** Benjamin Brachi, Danièle L. Filiault, Rahul Pisupati, Tal Dahan-Meir, Anna Igolkina, Alison Anastasio, Mathew S. Box, Susan Duncan, Talia L. Karasov, Envel Kerdaffrec, Laura Merwin, Timothy C. Morton, Viktoria Nizhynska, Polina Yu. Novikova, Fernando Rabanal, Takashi Tsuchimatsu, Torbjörn Säll, Caroline Dean, Svante Holm, Joy Bergelson, Magnus Nordborg

## Abstract

Local adaptation has been demonstrated in many organisms, but the traits involved, and the temporal and spatial scales at which selection acts are generally unknown. We carried out a multi-year study of 200 accessions (natural inbred lines) of Swedish *Arabidopsis thaliana* using local field sites and a combination of common-garden experiments that measured adult survival and fecundity, and selection experiments that measured fitness over the full life cycle. We found evidence of strong and variable selection, with particular genotypes favored more than five-fold in certain years and locations. Fecundity showed evidence of classical local adaptation, with accessions generally performing better closer to their home. However, relative to other accessions, southern ones usually had the highest fecundity—but were far more sensitive to harsh winters and slug herbivory, which strongly decreased both survival and fecundity. Furthermore, accessions originally sampled on beaches had low fecundity in all common-garden experiments, but massively outperformed all other accessions in the selection experiments, presumably due to an advantage during seedling establishment associated with their very large seeds. We conclude that local adaptation in *A. thaliana* reflects strong temporally and spatially varying selection on multiple traits, generally involving trade-offs and different life-history strategies, making fitness difficult to predict and measure.

## Introduction

Species reside in a heterogeneous landscape, with selective forces varying across time and space. When these selective forces are sufficiently consistent and spatially restricted, they may result in local adaptation whereby genotypes perform better at home than away (reviewed in *Savolainen et al., 2013*). The first demonstrations of such adaptive differentiation among populations focused on plants, due in part to the ease of reciprocal transplant "common-garden" experiments (***Turesson, 1922***; ***Clausen et al., 1940***). This tradition has continued in plant biology until the present day (***Sork, 2017***). Reciprocal transplant studies typically involve either phenotypically differentiated ecotypes (***Turesson, 1922***), presumably resulting from strong selection, or geographically disparate accessions that have a long history of evolving independently. But selection plays out in a dynamical landscape with changing selective pressures, interbreeding genotypes, and genetic variation that is restricted to the regional pool. Within this more realistic context, we know very little about the relevant spatial and temporal scales at which selection operates, and how short-term selection on various traits combine across time and space to shape long-term patterns of evolution.

At the genetic level, the picture is similarly murky, due in part to issues of scale. Even seemingly homogeneous fields vary on a micro-scale, with impacts on the relative fitness of plant genotypes (***Antonovics et al., 1987***; ***Stratton, 1994***, ***1995***). Such heterogeneity forms a mosaic that jointly determines evolution locally; for example, ***Frachon et al.*** (***2017***) studied rapid evolution in a natural population of *A. thaliana* and found that SNPs associated with intermediate levels of pleiotropy, not only across traits but across micro-environments, were those that changed in frequency. Results like these suggest a rich complexity in how short-term selection combines across time and space.

The present study sought to gain insight into local adaptation in *A. thaliana*, a widely distributed, highly selfing, mostly winter-annual plant that requires disturbed habitats and is consequently frequently found as a human commensal. While not the first study on local adaptation in *A. thaliana*, past work, including our own, had involved accessions collected across broad geographic ranges (***Hancock et al., 2011***; ***Fournier-Level et al., 2011***; ***Ågren and Schemske, 2012***; ***Ågren et al., 2013***)—and we were concerned that fitness differences observed on this scale would either be dominated by well-known phenomena, such as the connection between flowering time and latitude, or be too large (*i.e.*, the mortality of transplants would be too high) to reveal mechanisms. We were also worried that the traditional common-garden experiments used in most studies did not consider selection during seedling establishment, which we suspected would play a major role in an annual, weedy species like *A. thaliana*.

For these reasons, we decided to carry out a study in Sweden, using 200 Swedish accessions, most of which we had collected ourselves and whose habitats we were therefore familiar with (***Figure 1***). The study was focused on differences between the High Coast region in northern Sweden, where *A. thaliana* is mostly found on naturally eroded, south-facing slopes, and the heavily agricultural Skåne region in southern Sweden, where the species can be found in a variety of habitats, most of them associated with human disturbance, although it is also common on naturally disturbed beaches. In an attempt to include seedling establishment, we complemented standard common-garden experiments with novel selection experiments in which we sowed equal numbers of seeds of each accession in suitable natural habitats hoping to estimate fitness as presence in later generations—although it is important to note that when the study was started 15 years ago, we had no idea when it would be economically feasible to genotype any survivors.

**Figure 1.**
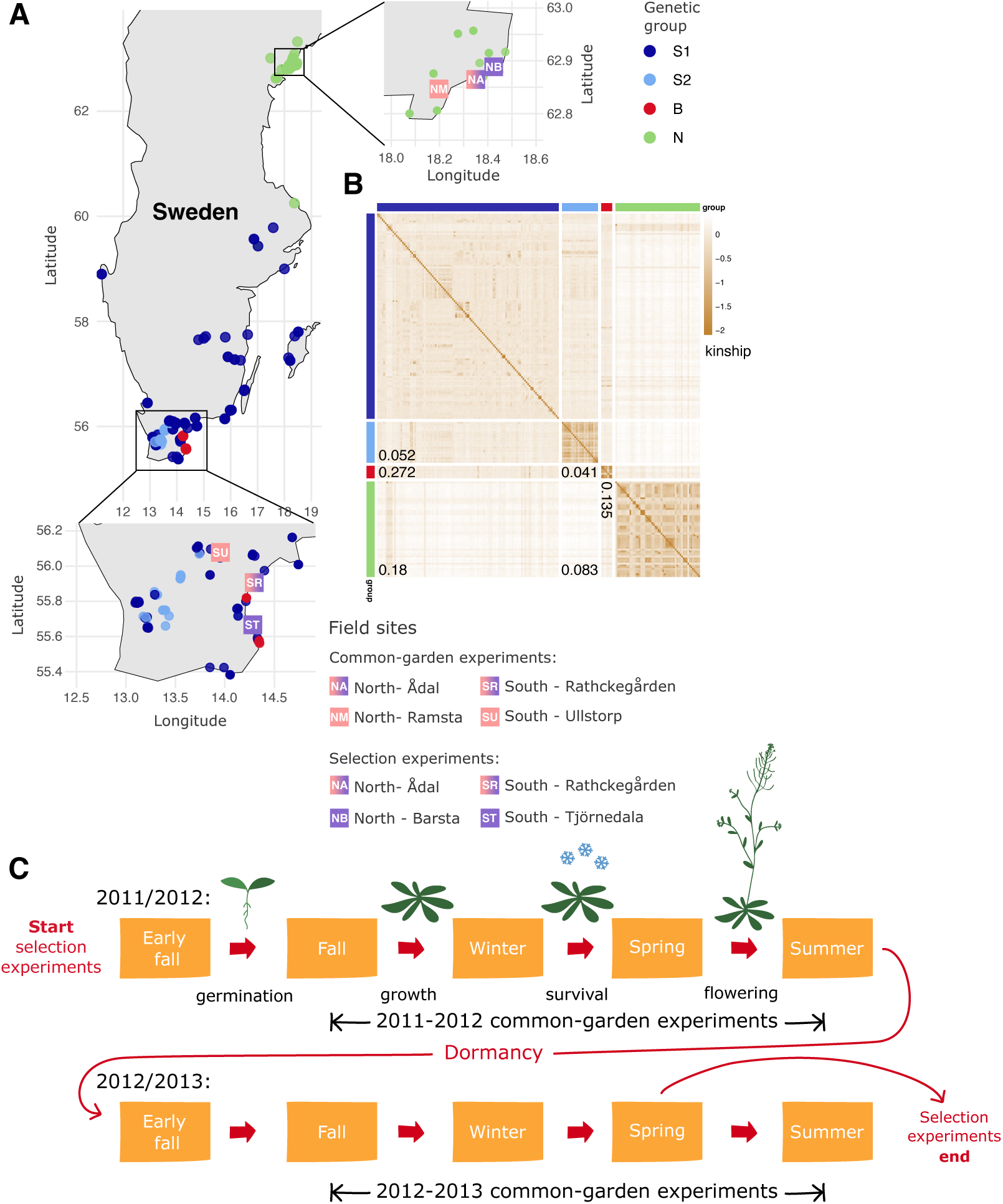
Experimental setup. (A) Location of the six field sites and origin of the 200 accessions. Note that sites NA and SR were used in both types of experiments. (B) Heatmap showing kinship between the accessions, hierarchically clustered by similarity. Marginal colors indicate membership in one of four genetic groups—the same colors are used for the sampling locations in panel A. Numbers are *F_ST_* estimates between groups. (C) Schematic of the experiments. In the common-garden experiments, which were replicated over two consecutive seasons, established seedlings were transplanted into the field in fall (using trays with holes in the bottom so that plants would root in native soil), and overwinter survival and fecundity assessed the following spring. In the evolution experiments, seeds were sown directly on the ground in equal numbers in early fall 2011 and surviving descendants sequenced before flowering in 2013. **Figure 1**—figure supplement 1. Population structure in the 200 Swedish accessions. **Figure 1**—figure supplement 2. The correlation of allele frequencies between the genetic groups. **Figure 1**—figure supplement 3. Photos of field sites: common gardens **Figure 1**—figure supplement 4. Photos of field sites: selection experiments in the south **Figure 1**—figure supplement 5. Photos of field sites: selection experiments in the north **Figure 1**—source data 1. Table of accessions used. **Figure 1**—source data 2. SNP matrix.

## Results

### Characterizing population structure

Although preliminary data had suggested clear genetic differentiation between northern and southern Sweden (***Nordborg et al., 2005***; ***Atwell et al., 2010***), the full extent did not become clear until genome-wide SNP data from larger numbers of accessions became available (***Long et al., 2013***; ***Huber et al., 2014***; ***1001 Genomes Consortium, 2016***). To describe the structure of the sample used here, we let ADMIXTURE (***Alexander et al., 2009***) divide the 200 accessions into distinct (supposedly randommating) groups. As with all clustering methods, the interpretation is quite arbitrary: we found that four groups provided a good fit to the geographic origin of the accessions and the overall structure of the sampling (***Figure 1***, ***Figure 1***—***figure Supplement 1***). 54 northern accessions fell into one group, N; S2 consists of 23 accessions exclusively from Skåne (Sweden’s southern-most province), with 60% being collected in urban areas; and B consists solely of 7 accessions collected directly on Baltic Sea beaches in southern Sweden. The more heterogeneous S1 group (116 accessions) is predominantly southern, but includes many isolated accessions from the rest of Sweden. The divergence between these groups can be seen from the kinship matrix, the *F*_ST_ values (***Figure 1***), and also the correlation in SNP allele frequency between groups (***Figure 1***—***figure Supplement 2***).

### Survival conditional on establishment in common gardens

Overwinter survival varied greatly between sites and years. In 2012-13, overall mortality was less than 2% in all four sites, and there was effectively no meaningful variation for survival (***Figure 2***). The same was true for one of the southern sites in 2011-12, SU, but the remaining three sites saw enough mortality to allow us to compare accessions and groups.

**Figure 2.**
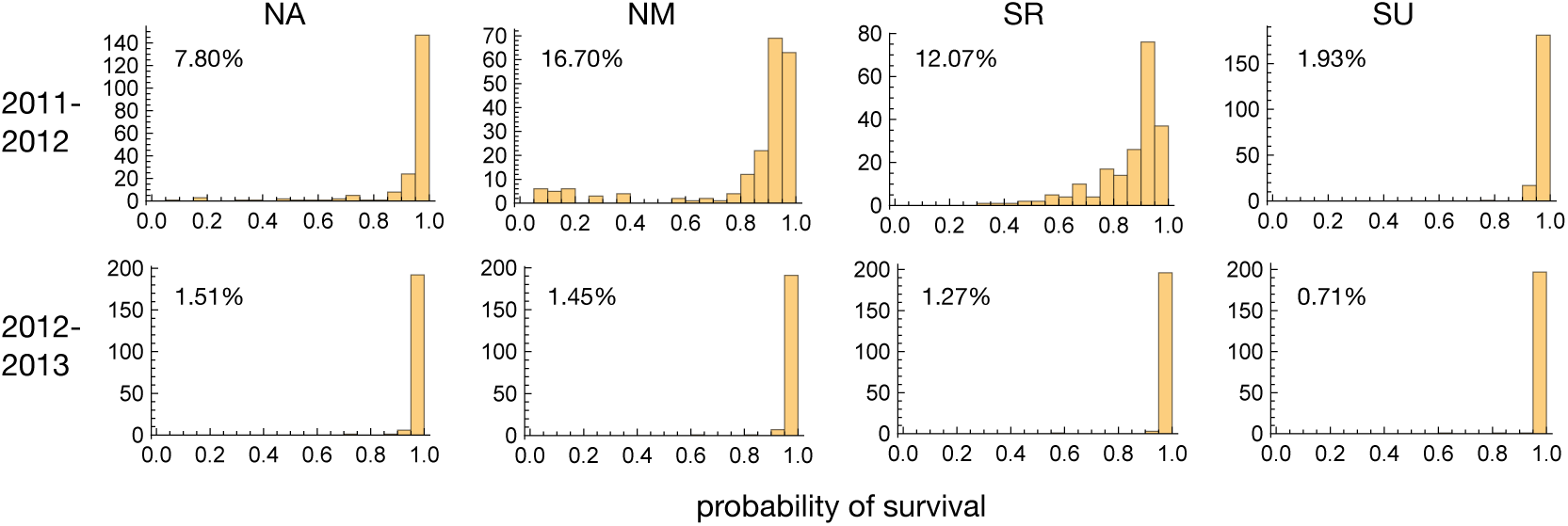
The distribution of accession survival probabilities across sites and years. Numbers in plots give total mortality in each experiment. **Figure 2**—figure supplement 1. The distribution of survival probabilities across experiments by group. **Figure 2**—source data 1. Overwinter survival data.

In these three experiments, southern accessions experienced dramatically increased mortality. In the two northern sites, NA and NM, where mortality was over 50% for many accessions, members of the S1 group were particularly affected, and our results suggest that conditions were more severe in NM both because overall mortality was twice as high (***Figure 2***) and because accessions with high mortality in NA always had high mortality in NM, whereas the converse was not true (***Figure 3***). Supporting this, a standard liability-threshold model (see Methods) with higher exposure in NM fits the data well.

**Figure 3.**
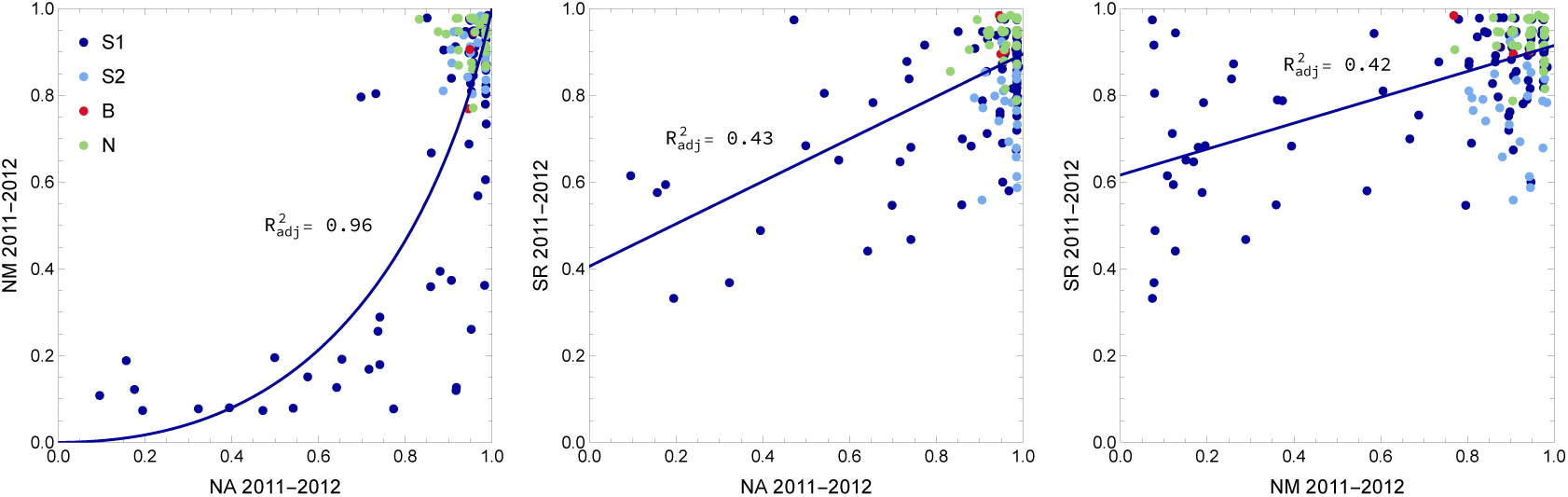
The joint distribution of accession survival probabilities between the three experiments with significant mortality. The curves were fitted using the S1 accessions only.

Occasionally reduced overwinter survival of southern accessions in northern sites is not surprising and suggests lack of adaptation to northern winters. It is consistent with the results of ***Oakley et al.*** (***2023***), who, using a cross between an Italian and a northern Swedish accession (***Ågren et al., 2013***), observed that Italian genotypes survived northern winters less well than northern Swedish genotypes in some, presumably harsher, years—whereas there was no difference in milder years.

In contrast, higher mortality of southern accessions in the southern SR site in 2011-12 was not expected. While only S1 accessions experienced higher mortality in the two northern sites, S2 accessions were also strongly affected in the SR experiment, suggesting a different cause of mortality (***Figure 3***, ***Figure 2***—***figure Supplement 1***). As it turns out, a major cause was observed: herbivory. The 2011-12 SR experiment was attacked by slugs, which caused substantial damage to leaves in the fall. We scored this damage and found that it both predicted overwinter survival and varied substantially between accessions, with S1 and S2 accessions being far more susceptible than B and or N accessions (***Figure 4***). Interestingly, despite the different causes, mortality in the SR experiment was correlated with mortality in the north for S1 accessions (***Figure 3***). In other words, the same accessions tended to die in both. This could reflect a direct causal connection, *e.g.* slug resistance also protects against cold. However, it seems more likely that some accessions are simply more sensitive to stress, be it from cold or slugs. Supporting this notion, survival in the SR experiment remained correlated with survival in the northern experiments after correcting for slug damage (***Figure 4***—***figure Supplement 1***).

**Figure 4.**
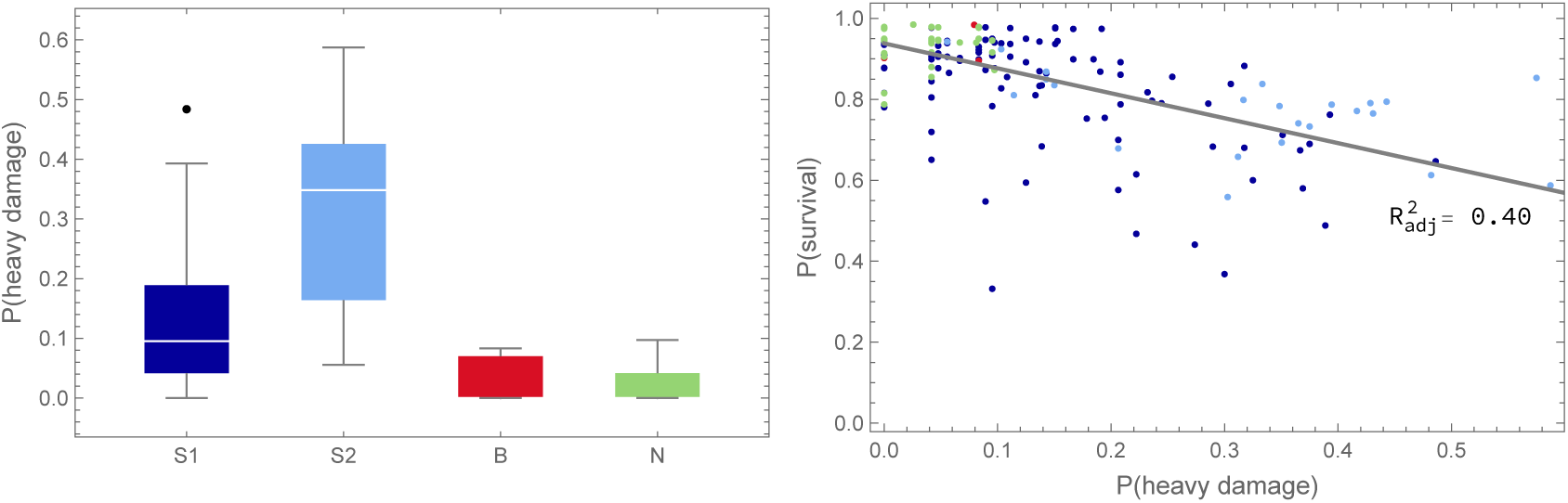
Overwinter survival was affected by herbivory. Left: Slug damage in the SR 2011-12 experiment affected groups differently (Kruskal-Wallis test: *p <* 0.01 for all comparisons). Right: Average slug damage for an accession decreased its probability of survival (ANOVA: *p* = 1.6 × 10^–23^). **Figure 4**—figure supplement 1. Overwinter survival was also affected by underlying susceptibility to stress. **Figure 4**—source data 1. Slug damage data.

GWAS for overwinter survival exhibit strong population structure confounding (***Figure 5***), which is precisely what is expected for traits that have been selected to differ between genetically differentiated groups (***Yu et al., 2006***; ***Zhao et al., 2007***; ***Atwell et al., 2010***; ***Platt et al., 2010b***). Standard statistical approaches for addressing this problem, like kinship-corrected mixed-linear models, have a tendency to over-correct, effectively "throwing out the baby with the bath water". In light of this, drawing conclusions from GWAS results alone, without supporting experiments, is not warranted. That said, we note that one of the top association peaks for survival in the SR 2011-12 experiment (***Figure 5***—***figure Supplement 3***) includes the *AOP* cluster, which has been shown to play a major role in natural variation for glucosinolate profiles and defense against herbivory, even on local scales (***Kliebenstein et al., 2001***; ***Gloss et al., 2022***). The peak appears to involve a haplotype over 30 kb in length, which could reflect the segregating inversion at this locus that has been reported to be strongly associated with glucosinolate profiles in Sweden (***Katz et al., 2021***). We confirmed that these profiles are strongly associated with the variation for slug damage (***Figure 6***), strongly suggesting a causal relationship. ***Katz et al.*** (***2021***) also report a major association at the *MAM* cluster, but the polymorphism responsible for this association does not segregate is Sweden, and hence it is not surprising that we find no association at this locus.

**Figure 5.**
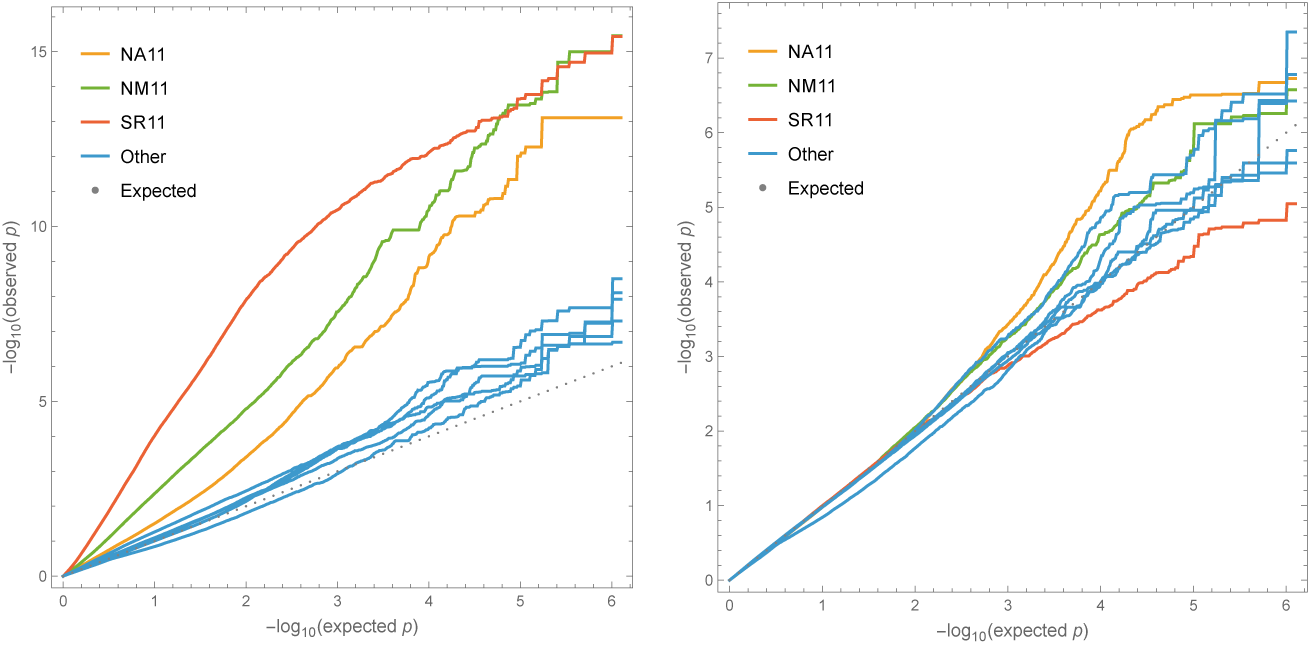
Quantile-quantile plots of p-values for overwinter survival GWAS against the expected uniform distribution. Left plot without correction for structure; right plot including a standard mixed-linear model kinship correction. Only SNPs with Minor Allele Frequency (MAF) greater than 5% were included to avoid outlier affects. Experiments with significant mortality exhibit genome-wide inflation of significance, while remaining experiments demonstrate that tails are inflated even for noise phenotypes. **Figure 5**—figure supplement 1. Manhattan plots without structure correction. **Figure 5**—figure supplement 2. Manhattan plots with structure correction. **Figure 5**—figure supplement 3. Zoom-in on *AOP* region. **Figure 5**—figure supplement 4. Zoom-in on *SVP* region.

**Figure 6.**
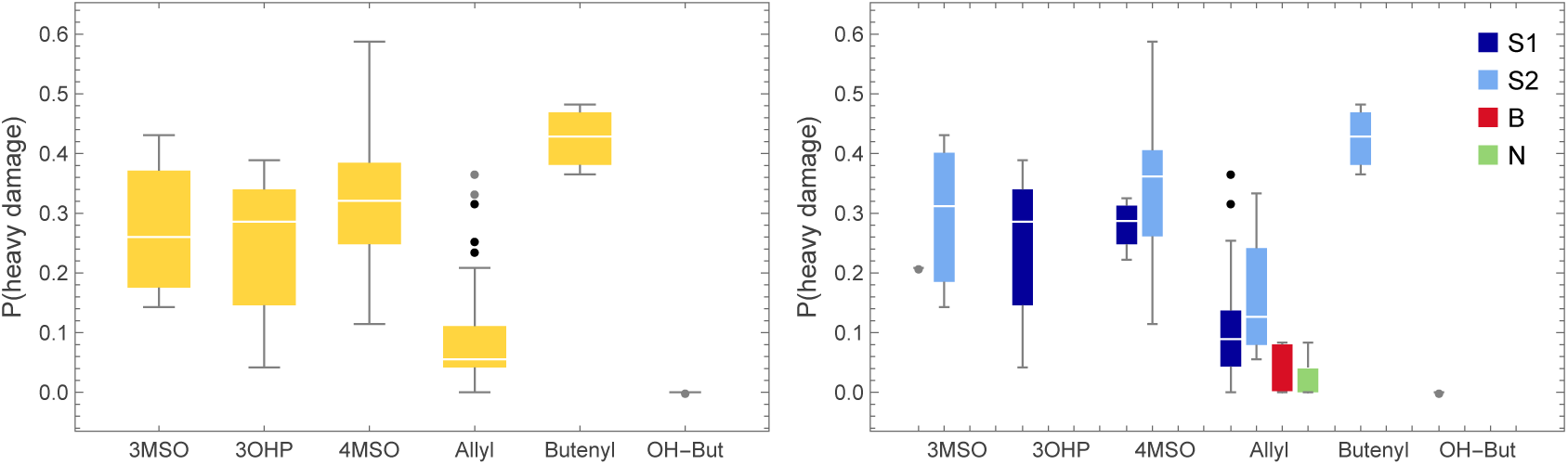
Slug damage was associated with glucosinolate profiles. Left: Slug damage in the SR 2011-12 experiment affected the glucosinolate profiles defined in ***Katz et al.*** (***2021***) differently (data for 122 of our accessions; Kruskal-Wallis test: *p <* 5 × 10^–13^). Right: The difference in mortality between groups (*cf.* Figure 4) is associated with glucosinolates profiles, with the putatively protective “allyl” phenotype present in 100% of B and N accessions, 79% of S1 accessions, but only 22% of S2 accessions. Figure 6—figure supplement 1. Survival was also associated with glucosinolate profiles.

In general, far less work has been done on overwinter survival than on traits like flowering time or glucosinolate profiles, and there are therefore fewer *a priori* candidates to help interpret GWAS results. A possible exception is *SVP*, which is under one of the strongest (uncorrected) association peaks in NM 2011-12 (***Figure 5***—***figure Supplement 4***). This gene encodes a MADS-box transcription factor that plays a major role in flowering-time variation (***Méndez-Vigo et al., 2013***; ***Guo et al., 2023***) via thermo-sensory pathways (***Lee et al., 2013***), and hence may also be involved in preparing plants for winter. *SVP* is frequently identified in selection scans because it exhibits unusually high geographic differentiation, including with latitude—consistent with a role in local adaptation (***Horton et al., 2012***; ***Zou et al., 2017***; ***Guo et al., 2023***). However, precisely because of this, *SVP* will be associated with any trait that shows similar differentiation and further experiments will be required to determine which adaptive traits it actually influences.

Interestingly, GWAS for overwinter survival in the north in 2011-12 finds no association at *CBF2*, which has been shown to play a role in freezing tolerance in the above-mentioned cross between a northern Swedish and an Italian accession due to a loss-of-function allele in the Italian parent (***Gehan et al., 2015***; ***Lee et al., 2024***). We used the BLAST-based simsearch tool from the Pannagram package (***Igolkina et al., 2026***) to look for the causal 13-bp deletion in close to 600 independently assembled *A. thaliana* genomes, including over 150 from Sweden (***The 1001G+ Consortium, 2024***), but found the deletion only in the Italian accession used as RIL parent by ***Ågren et al.*** (***2013***). This suggest that the loss-of-function allele identified by ***Gehan et al.*** (***2015***) is rare and does not play a major role in the genetic architecture of freezing tolerance in Sweden (or Europe-wide). Thus, although our results are consistent with those of ***Oakley et al.*** (***2023***) at the phenotypic level, the genetic basis appears to be different.

### Fecundity conditional on survival in common gardens

Fecundity was estimated from photos of harvested mature plants, thus also capturing the amount of biomass these annual plants invested in reproduction (see ***Brachi et al., 2022***, for details). Linear modeling of the entire study (*i.e.*, including all 8 experiments) showed that while the *accession*, *year*, and *site* terms were all highly significant, a greater proportion of explainable variation was due to interactions between them (***Figure 7***). Significant *year*site* and *year*site*accession* interactions demonstrate that fecundity varies from year to year, with this effect differing among sites. Importantly, interactions involving *accession* indicate that the genetic effect on fecundity varied between experiments—a pre-requisite for local adaptation.

**Figure 7.**
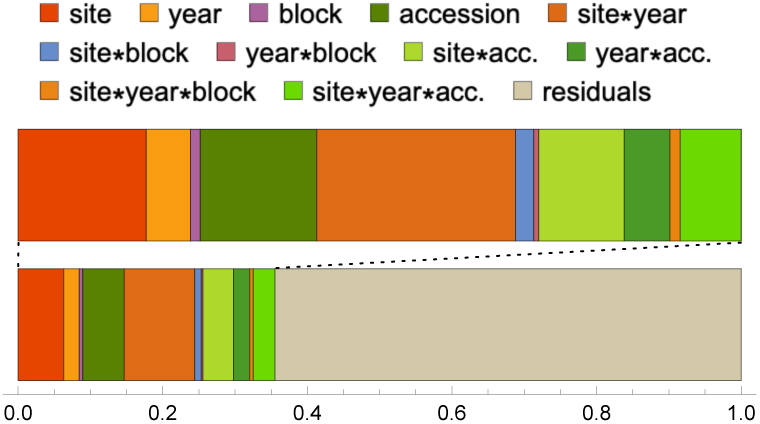
Variance-partitioning (ANOVA) from full model of fecundity. The top bar zooms in on the explained variance from the complete partitioning in the bottom bar. Figure 7—source data 1. Full ANOVA tables.

We investigated this further by considering the relative performance of the accessions (and, by extension, groups) in each of the 8 experiments. Briefly, we calculated Best Linear Unbiased Predictors (BLUPs) of the effect of accession in each experiment (see Methods) and used a standard Principal Components Analysis (***Figure 8***) to look for patterns across experiments. The first principal component (PC1; explaining 38% of the variance) identifies a common pattern across all experiments; the second component (PC2; 15% of the variance) switches sign between the years of the study; and the third (PC3; 13% of the variance) switches sign between northern and southern experiments—except that SR 2011-2012, the experiment with heavy overwinter mortality due to slugs, behaves like a northern experiment.

**Figure 8.**
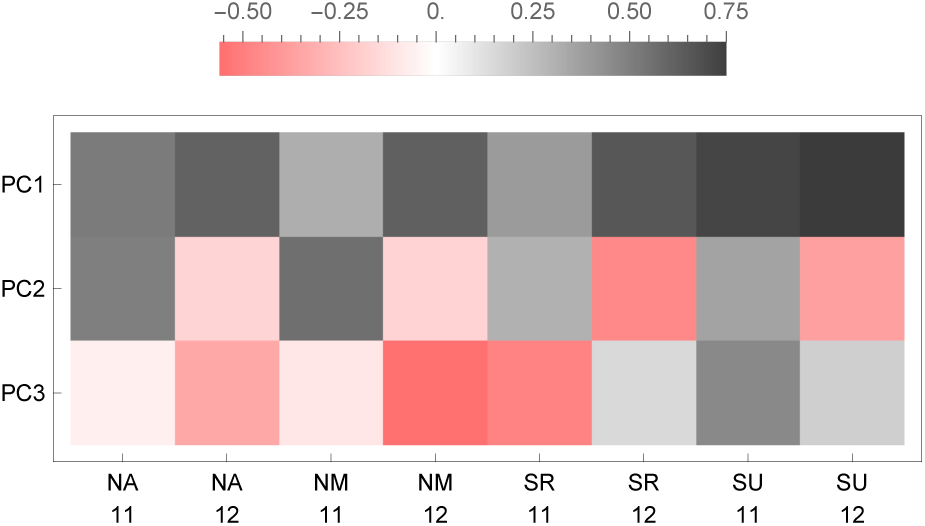
The loadings of the first three PCs from a PCA of the fecundity BLUPs on the eight experiments. **Figure 8**—figure supplement 1. The distribution of each PC by group. **Figure 8**—figure supplement 2. Positions of accessions in PC1-PC2 space. **Figure 8**—figure supplement 3. Positions of accessions in PC2-PC3 space.

The full distribution of BLUPs helps us make sense of these patterns by considering how the groups performed (***Figure 9***). PC1 reflects the fact that S1 and S2 accessions generally tend to have higher fecundity than B and N accessions. The B accessions have the lowest median fecundity in all 8 experiments (significantly so in only 5 out of 8, but note that the probability of any one group consistently being last is (1/4)^7^ = 6 × 10^–5^) and the N accessions have the second lowest fecundity in all southern experiments and all 2012-13 experiments (significantly lower than S1 and S2 in 4 out of 6 experiments). Only in the northern experiments in 2011-12 was this trend broken, although the differences are small and not statistically significant.

**Figure 9.**
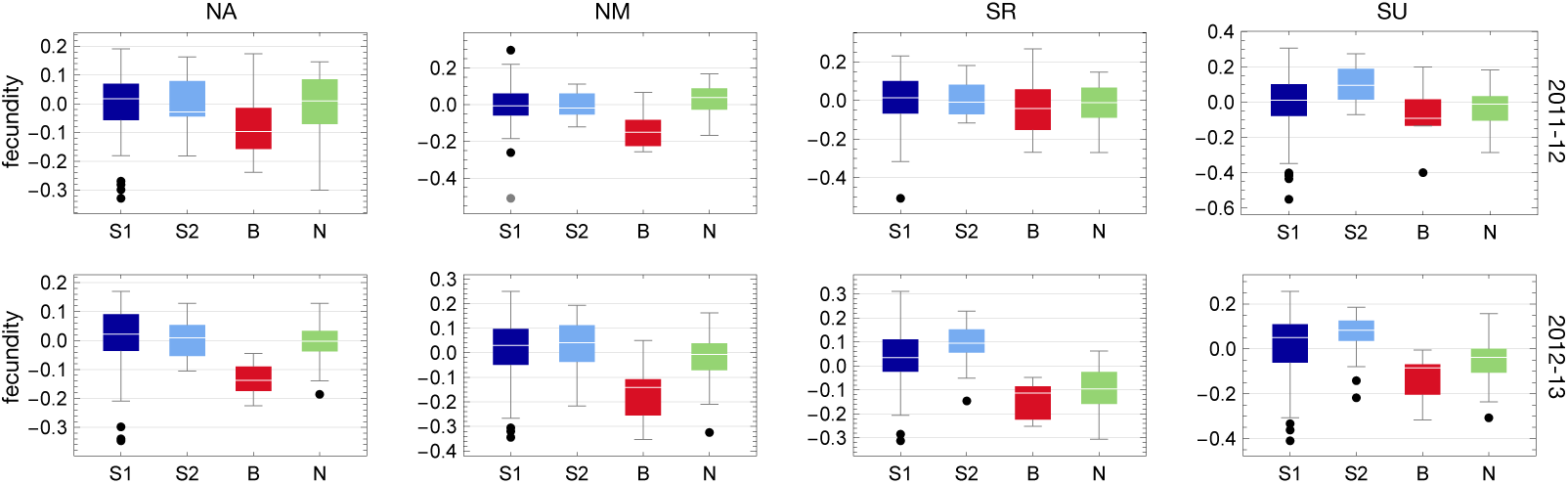
The distribution of fecundity BLUPs for each experiment by group. **Figure 9**—figure supplement 1. Accession fecundity is correlated with overwinter survival. **Figure 9**—source data 1. Fecundity BLUPs.

The pattern captured by PC1 is far stronger in 2012-13 than in 2011-12, and this is captured by PC2. Although the differences are sometimes small, S1 and S2 accessions generally had higher fecundity in 2012-13 than in 2011-12, while the reverse was true for B and N accessions (pattern holds in 13 out 16 comparisons, see ***Figure 10***).

**Figure 10.**
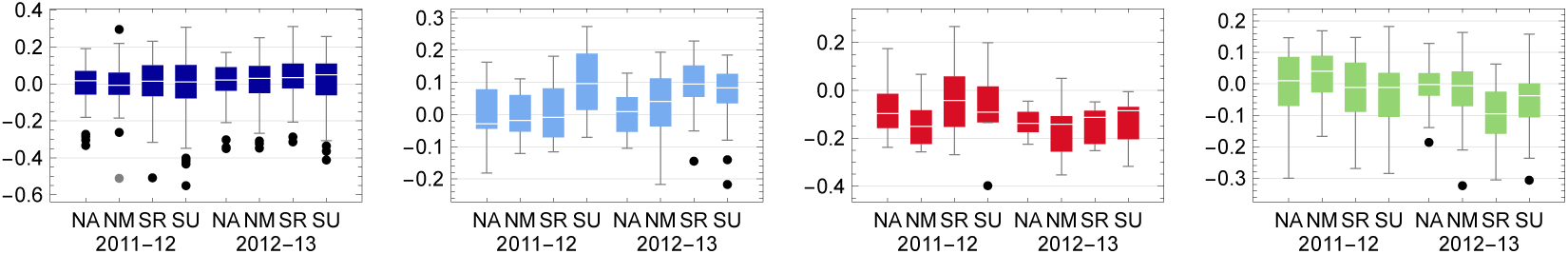
The distribution of fecundity BLUPs for each group by year and site (*cf.* Figure 9).

PC3, finally, captures the fact that, within years, southern accessions (S1, S2, and B) generally perform better in southern than in northern experiments, while the opposite is true for the northern accessions (N). The probability of this pattern in a particular experiment is 1/6, and it occurs in 7 of the 8 experiments (one-tailed probability = 2.4 × 10^–5^).

In the three experiments where high mortality was observed, fecundity was correlated with overwinter survival, suggesting that surviving members of affected genotypes were weakened by whatever caused death among their (inbred) siblings (***Figure 9***—***figure Supplement 1***). This effect thus decreased the fecundity of the S1 accessions in NA and NM 2011-12, and of the S1 and S2 accessions in SR 2011-12, presumably explaining why this latter experiment looks "northern" in the PCA (***Figure 8***).

In addition to fecundity, we also noticed variation in the extent to which leaves had turned purple in the fall, presumably as a sign of stress. We scored this from photos (see Rosette purpleness in Methods). Differences between accessions explained 4–26.6% of color variation across the six experiments in which it was called, and correlations with mortality and fecundity were mostly in-significant. Interestingly, having a purple rosette appeared to have a relatively simple genetic basis: in the experiment where accession explained the most variation, GWAS revealed single major locus in the anthocyanin production pathway, *PAP2* (***Figure 11***). The 10 top SNPs, all located between 24,764,623 and 24,783,821 bp on chromosome 1, jointly explained 17.7% of genetic variation for purpleness among accessions.

**Figure 11.**
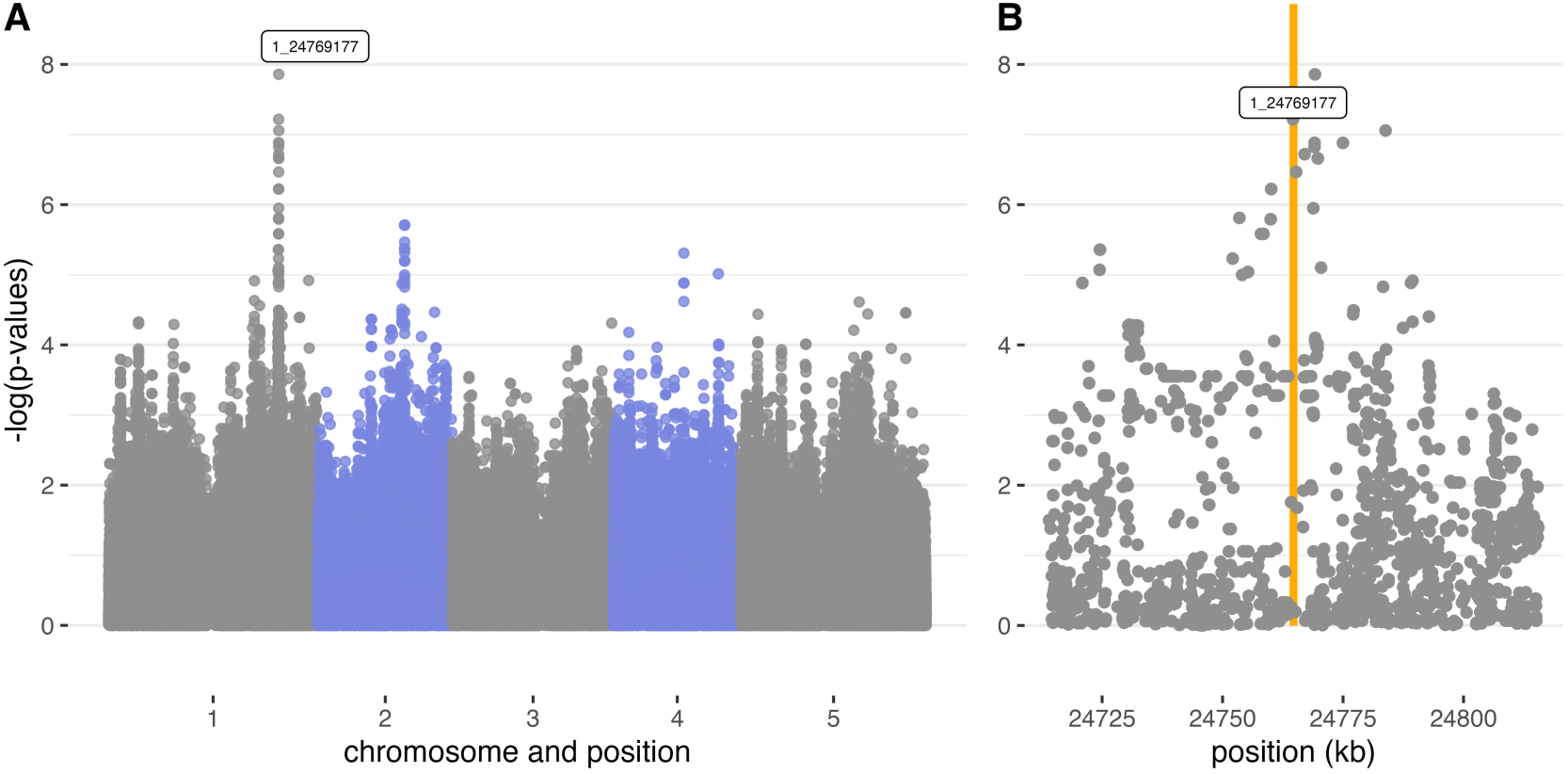
GWAS of rosette purpleness in SU 2011-12 identifying *Production of Anthocyanin Pigment 2, (PAP2, AT1G66390)*, located on chromosome 1 between 24,763,941 and 24,765,541 bp. A. Genomen-wide Manhattan plot B. Zoom-in on the 100 kb window around *PAP2*, represented by the orange bar.

In contrast, fecundity is obviously a complex trait. It is conditional on overwinter survival (in itself a complex trait), and it is characterized by complex genotype-by-environment interactions. In addition, it is strongly correlated with population structure. As expected, GWAS results thus exhibit strong population-structure confounding, and while attempting to eliminate this confounding statistically eliminates false positives, it also removes signal (***Figure 12***).

**Figure 12.**
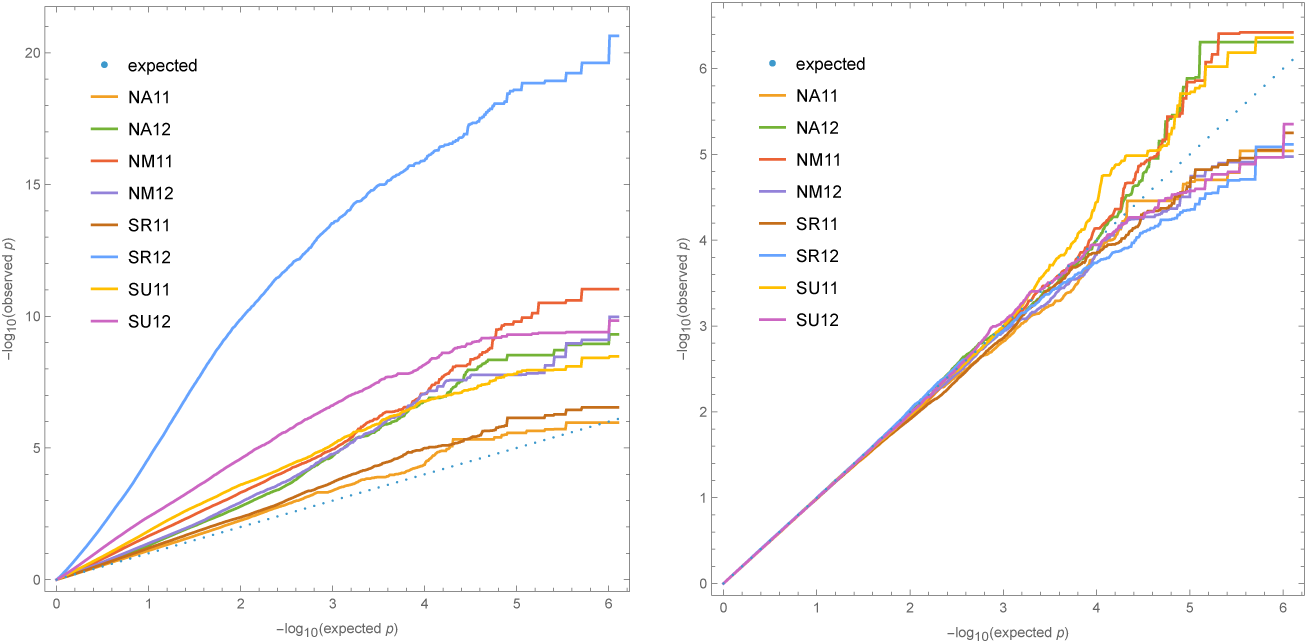
Quantile-quantile plots for GWAS fecundity data. Left plot without correction for structure; right plot including a standard mixed-linear model kinship correction. The extreme confounding for SR 12 reflects the unusually large difference between S1/S2 and B/N in this experiment (Figure 9). Only SNPs with MAF greater than 5% were included to avoid outlier affects. Figure 12—figure supplement 1. Manhattan plots for fecundity GWAS without correction for structure. Figure 12—figure supplement 2. Manhattan plots for fecundity GWAS with correction for structure.

That said, the GWAS identify many clear peaks of association that are suggestive of large-effect polymorphisms. Some of these are shared between experiments, consistent with the patterns discussed above (*e.g.*, the dominant S1/S2 vs B/N patterns identified by PC1 in ***Figure 8***). Each peak includes on the order of ten genes, but in the absence of strong priors for which genes might affect a highly complex trait like fecundity, speculating about causality is futile. These results will hopefully guide future experiments.

### Fitness in selection experiments

#### Estimating fitness

Our experimental evolution experiments were also carried out at four sites, two in the north, and two in the south, with one site in each region associated with a common garden experiment (***Figure 1***). One of the southern sites, ST, was located on a beach, with members of the B group as its local population. Although native plants were growing within less than a hundred meters, the sites were assessed to be free of *A. thaliana*, but were deemed to be suitable habitats (see photos in ***Figure 1***—***figure Supplement 4*** and ***Figure 1***—***figure Supplement 5***). We were correct about the second part, but not the first, as we shall see below.

At each site, seeds from the 200 accessions were mixed in equal numbers and sown at relatively high densities to establish three to five independent experimental plots, which were then allowed to complete a full life cycle under natural conditions (including seed dormancy, seedling establishment, and competition). At least 70 plants per plot (1,238 plants in total) were randomly sampled after the second winter. These plants were individually genotyped using low-coverage short-read sequencing to assess the composition of each experimental population. Given that all accessions started the experiment at equal frequency, fitness for each accession in each site was simply calculated using its sampled population frequency at that site.

Genotyping turned out to be challenging, mostly because of the difficulties involved in extracting DNA from plants that were collected under field conditions and which were often tiny and half-dead. As described in Methods (see ***Figure 20***), we concluded that, of 1,174 genotyped samples: 75.4% were homozygous individuals matching our 200 experimental accessions, 15.8% were homozygous individuals that did not match our experimental accessions (*i.e.*, they were native “volunteers"), and 8.8% were extensively heterozygous (either due to recent outcrossing or sample contamination). The native volunteers were found in every experiment (***Figure 20***—***figure Supplement 1***)—thus demonstrating that we did indeed pick sites suitable for *A. thaliana*—and they do not pose a problem, as they do not affect the relative fitness estimates for the experimental accessions.

However, the presence of *bona fide* natives alerted us to the fact that our fitness estimates could be confounded by the inclusion of natives indistinguishable from our experimental accessions. To investigate this we used background knowledge of the population structure in *A. thaliana* to identify experimental accessions that had originally been collected close enough to an experimental site for it to be plausible that nearly identical plants could be native to that site (see Methods). We found that this problem could potentially be serious in two of the sites: NB, where 22% of putatively experimental samples were potentially natives, and ST, where the proportion of potential natives was 52%. The proportion of potential natives at NA was only 3% and the SR site could not have been affected as no experimental accession had been collected nearby.

As it turns out, two accessions make up most of these potential natives and are also obvious outliers in the expected experimental populations (***Figure 13***). At the NB (Barsta) site, the accession Bar1 (originally sampled in Barsta) comprised 20% of sampled individuals (and 88% of potential natives). While it is theoretically possible that Bar1 is extremely well-adapted to its local habitat and thus outperformed all other accessions, increasing 40-fold in frequency during the course of the experiment (all accessions started the experiment at frequency 0.5%), it seems far more likely that most of the Bar1 individuals sampled at NB hail from the native, non-experimental population.

**Figure 13.**
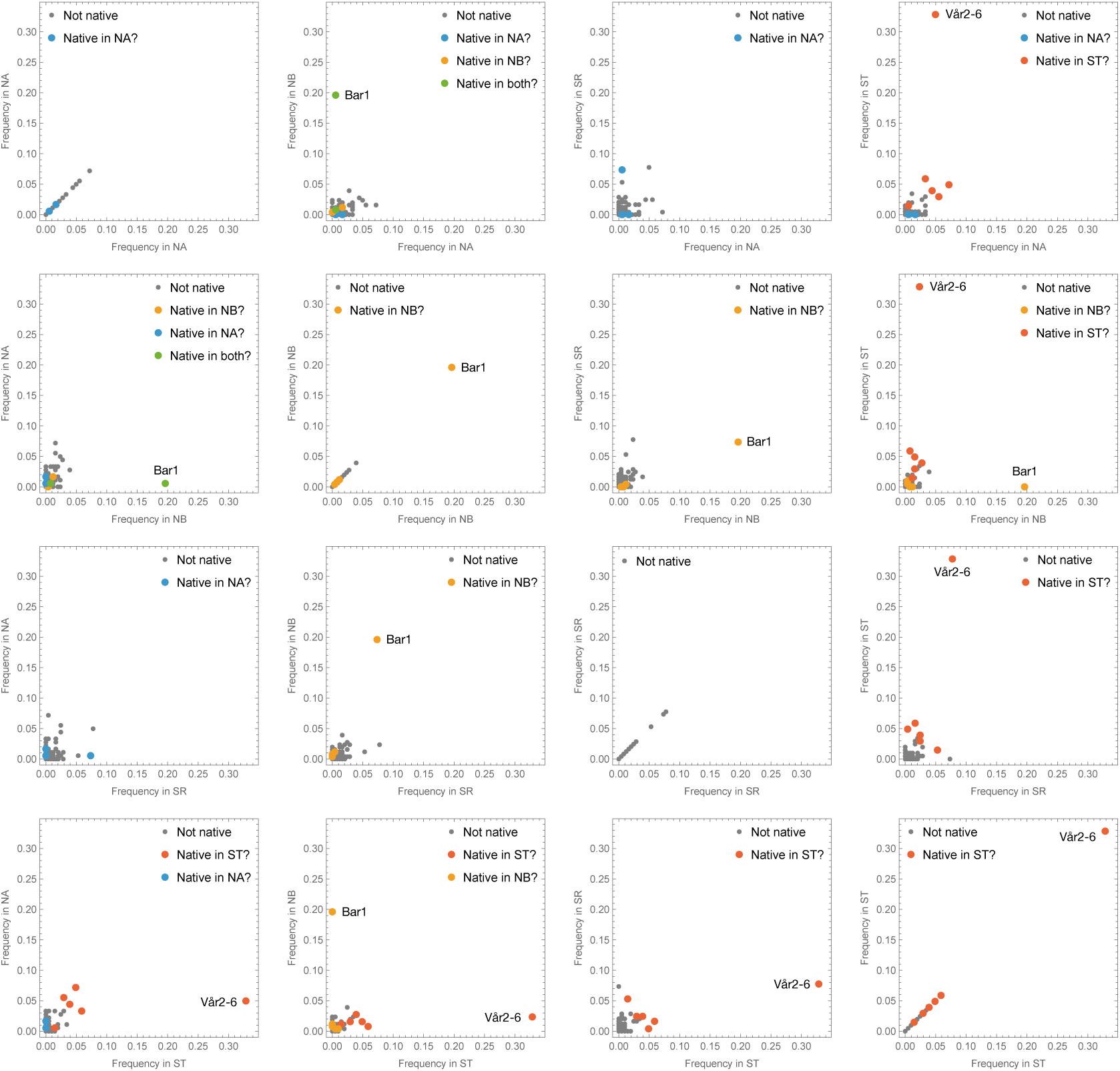
Scatter-plots comparing estimated accession frequencies in the four selection experiments. Accessions that are potential natives in a site are indicated by color and the two obvious outliers (Bar1 in NB, and Vår2-6 in ST) are highlighted. Figure 13—figure supplement 1. Same as Figure 13, but with the two obvious outliers removed to show remaining data better. Figure 13—source data 1. Estimated fitnesses from evolution experiments.

Similarly, at the ST (beach) site, the accession Vår2-6, originally collected about 3.5 km further south along the same beach, comprised 33% of sampled individuals, an implied 65-fold increased in frequency. Removing these two accessions from the study eliminates 69% of all potential natives, effectively eliminating the problem at all sites except for ST, which still has 28% potential natives. This reflects the facts that all experimental accessions hailing from the beach with the ST site are potential natives, *and* that these accessions were over-represented in the individuals sampled in the ST experiment—*prima facie* consistent both with higher fitness of beach accessions and confounding by the native, non-experimental population. However, looking across experimental sites (***Figure 13***, ***Figure 13***—***figure Supplement 1***), we see that these accessions were over-represented in *all* experiments, including at the three sites where their numbers could not have been inflated by native contributions, and we therefore conclude that they actually had higher fitness and treat them as experimental samples. As we shall see below, other observations support this conclusion. Nevertheless, in the ST site, the fitness estimates maybe be inflated by the inclusion of natives, but none of our conclusions depend on this.

Importantly, the eliminated beach accession, Vår2-6, also had very high fitness in the other experiments (5.0% in NA, 2.4% in NB, 7.8% in SR) suggesting that its extraordinarily high estimated fitness in its native habitat (ST) probably was not *only* due to confounding by natives, but also reflect general fitness advantage plus local adaptation. Excluding this accession thus probably biases the fitness of the B group downwards. Note that the same is not true for the Bar1 accession (0.6% in NA [where it was also potentially native], 7.3% in SR, 0% in ST).

#### Comparing accessions and groups

The data presented in the previous section already reveal the most striking conclusion from our selection experiments, namely that the B accessions tended to outperform all other accessions in all sites (***Figure 14***). This unexpected result contrasts sharply with the observation that the B accessions had the lowest fecundity in all common-garden experiments (***Figure 9***), and obviously does not support any simple notion of local adaptation. When considering what might provide these accessions with up to a 20-fold fitness-advantage (up to 25 in ST, although this estimate may be inflated), we remembered that some of them tend to produce very large seeds, which might be an advantage in seedling establishment, if for no other reason than that seedlings from large seeds grow faster (***Slovak et al., 2020***; ***Clauw et al., 2022***).

**Figure 14.**
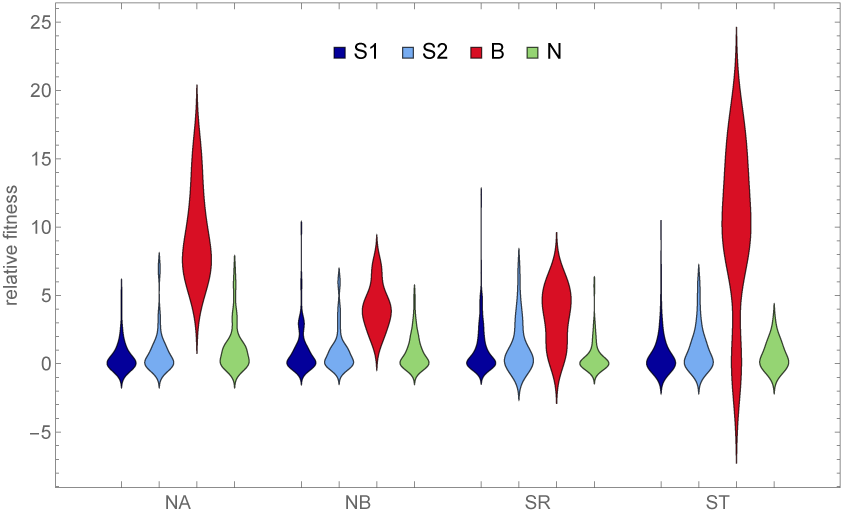
The distribution of the estimated relative fitness by genetic group (same data as in Figure 13 but rescaled by dividing by the expected frequency in the absence of selection).

Using existing seed size measurements from different experiments, we discovered that our genetic groups did indeed differ dramatically in seed size, with the B accessions producing by far the largest seed (***Figure 15***). Importantly, these differences appear to be insensitive to environmental conditions (***Figure 15***—***figure Supplement 1***). Plotting fitness against seed size confirms that there is a relationship—but it is not a simple correlation. In particular, while N accessions also have large seeds (albeit not as large as B accessions), they have no higher fitness than S1 and S2 accessions. Large seed size seems to be necessary but not sufficient for high fitness.

**Figure 15.**
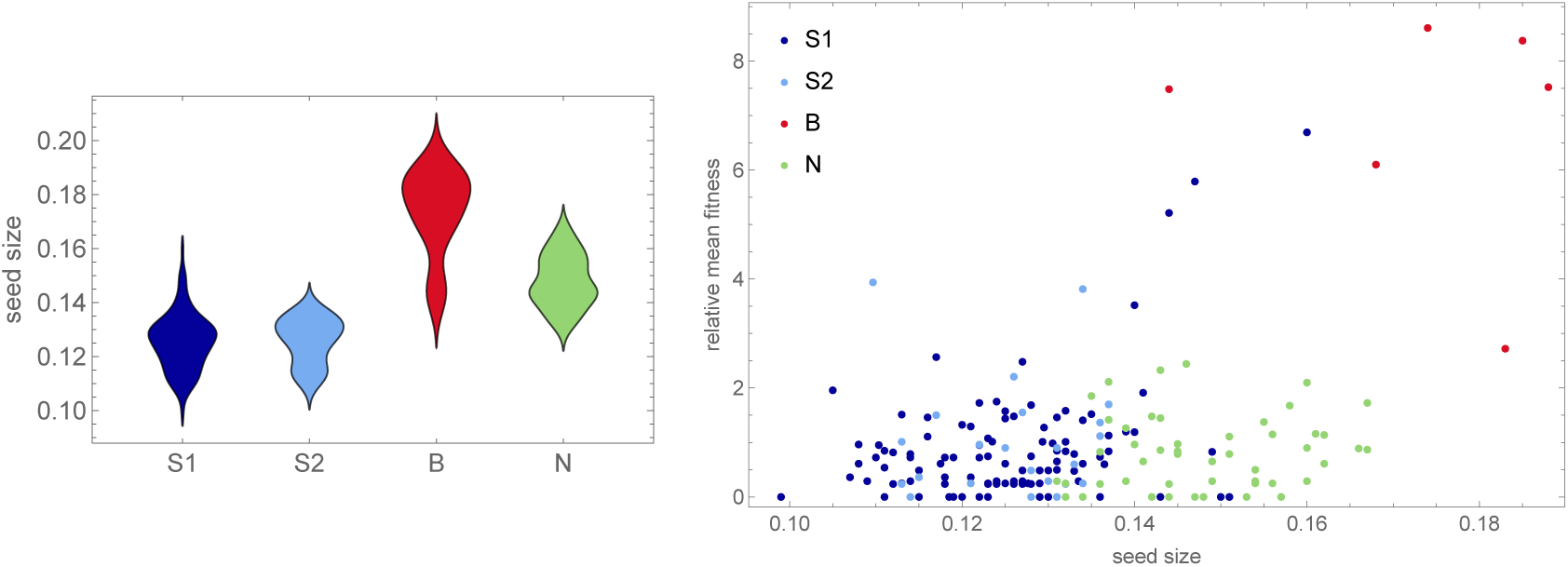
The distribution of seed size (data from ***Clauw et al., 2022***) by genetic group, and, on the right, mean fitness (across sites, from Figure 14) as a function of seed size. Figure 15—figure supplement 1. Correlation of seed size across environments. Figure 15—source data 1. Seed size estimates for project accessions.

Inspired by these results, we examined other traits that might contribute to fitness variation, in particular those related to seedling establishment. Based on field observations, we knew that B accessions tended to have very long roots—likely a necessity for growing on sandy beaches. ***Slovak et al.*** (***2020***) measured early root growth experimentally and showed that it depends on seed size, but also that it has a genetic basis that appears to be independent of seed size (***Slovak et al., 2020***). Their studies include a small subset of the Swedish accessions used here, and indeed we see that the one high-fitness B accession included is an outlier for rapid root growth when seed size is not accounted for, and remains at the top even when it is (***Figure 16***, panels A–B).

**Figure 16.**
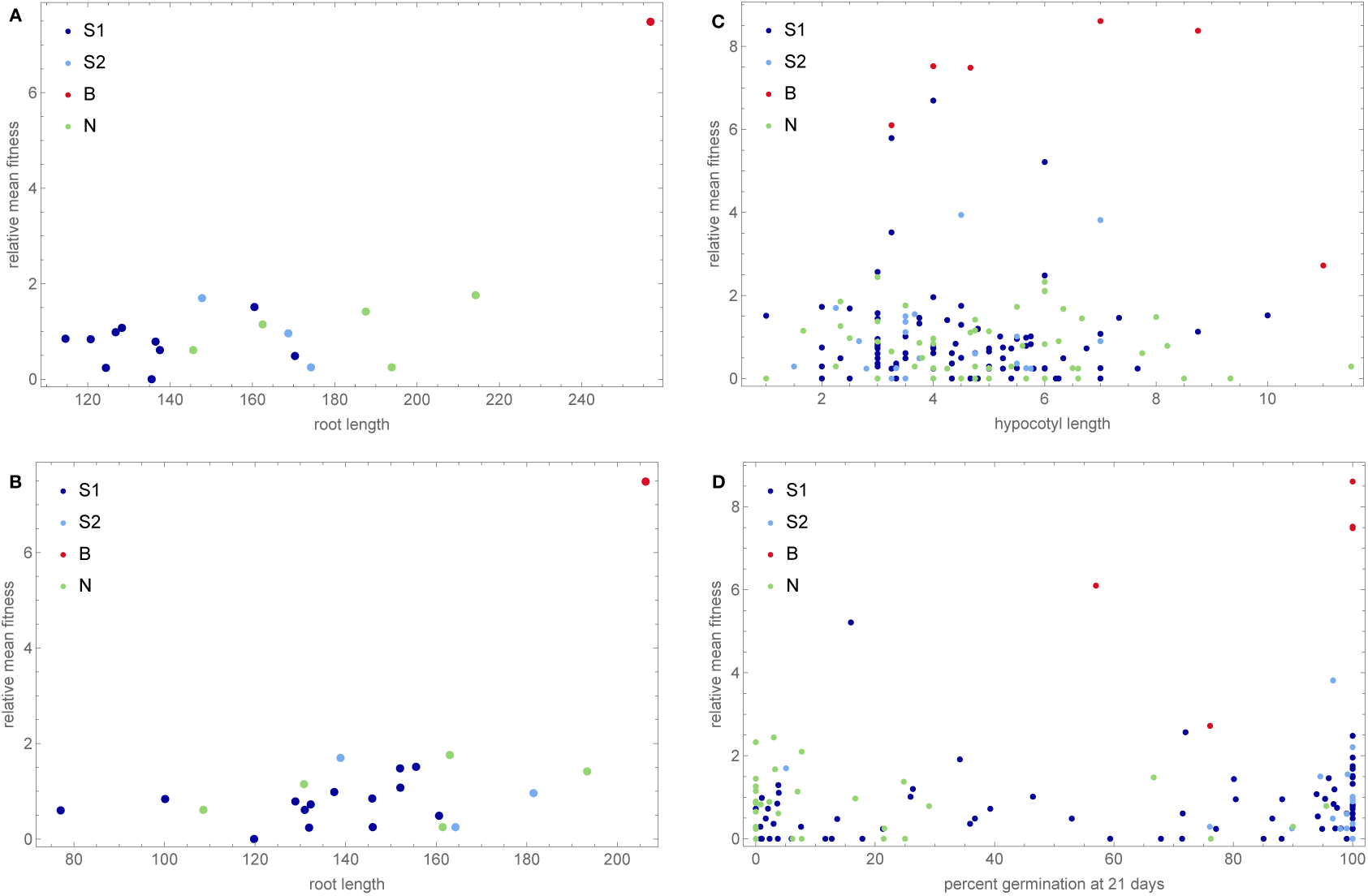
Mean fitness as functions of various phenotypes related to seedling establishment. A. Root length, from ***Slovak et al.*** (***2020***). B. Same as A, but corrected for seed size, which was positively correlated with early root growth. C. Hypocotyl elongation. D. Primary seed dormancy, from ***Kerdaffrec et al.*** (***2016***). Figure 16—figure supplement 1. Fitness as a function of hypocotyl elongation in each experiment. Figure 16—figure supplement 2. Fitness as a function of primary seed dormancy in each experiment.

Another plausible trait is hypocotyl elongation, which could be especially useful for buried seeds. We tested this experimentally, but no obvious pattern was found (***Figure 16***, panel C). Finally, we had already shown that B accessions tended to have high primary dormancy, whereas N accessions have very low primary dormancy (***Kerdaffrec et al., 2016***). We interpreted this as local adaptation to hot and dry summers on southern beaches vs adaptation to very short growing season in the north. Consistent with this, there is no obvious relationship between seed dormancy and mean fitness across our experiments (***Figure 16***, panel D), but the pattern seen in individual experiments does support dormancy playing a role in local adaptation (***Figure 16***—***figure Supplement 2***).

Note also that, while the B accessions stand out as having very high fitness, fitness is not perfectly correlated with genetic group. In addition to five B accessions, the top right corner of the scatter plot in ***Figure 15*** contains three S1 accessions that have lower fitness and seed size, but are still positive outliers. Two of these hail from the same beaches as the B accessions and the third had been sampled from another sandy beach—on the island of Gotland in the middle of the Baltic Sea, over 500 km away (***Figure 1***). To confirm that the apparent association between seed size and beach habitat is real, we measured seed size in 246 additional Swedish accessions. The pattern was confirmed (***Figure 17***).

**Figure 17.**
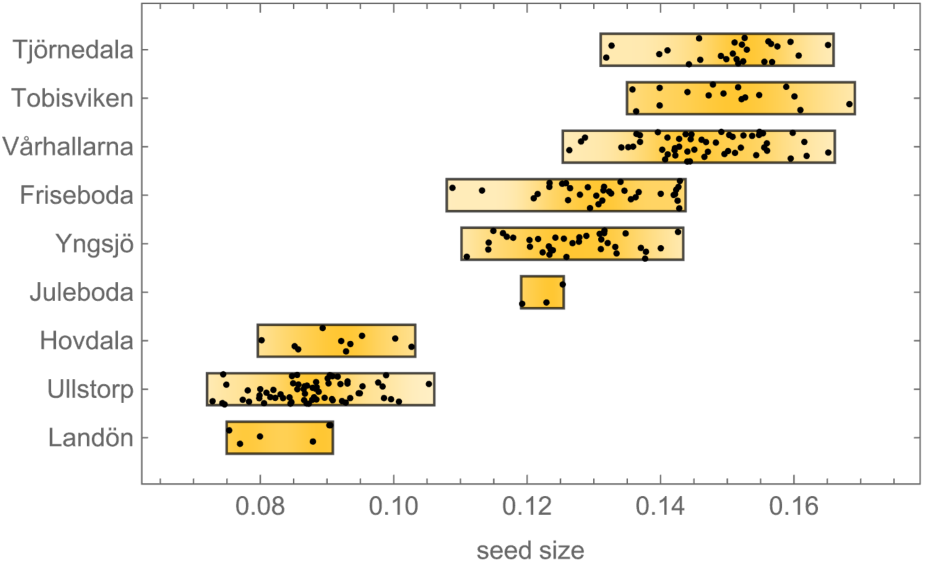
Seed size distribution for 246 additional accessions sampled from 9 sites in southern Sweden in 2017 (see Methods). The top 6 sites were located on beaches and the bottom 3 were located inland. The experimental B accessions hail from the top four sites. Figure 17—source data 1. Seed sizes for these accessions.

#### Estimating allele-frequency changes

To gain insight into the genetic basis of fitness, we considered changes in SNP frequencies during the course of the experiment. We used basic population genetics to model changes in the absence of selection, and estimated significance using standard statistics (see Methods). Because the results across experiments were highly correlated, we combined results by simply multiplying *p*-values. The results of this selection scan can be visualized like GWAS results using a Manhattan plot, and look very similar. They are also plagued by population structure—as for GWAS results, we see horizontal bands of identical p-values reflecting extensive haplotype structure and long-range linkage disequilibrium (***Figure 18***). Another manifestation of population structure confounding is the curious relationship between p-values from a seed size GWAS and from the selection scan. As can be seen in the right panel of ***Figure 18***, this distribution has two main axes. The lower one captures the total variation in seed size, which is associated with S1 and S2 vs N and B, but is not strongly associated with fitness—N accessions do not have high fitness despite their larger seeds (***Figure 15***). The upper one captures only the component of seed size variation that is also associated with fitness, *i.e.*, B accessions vs everyone else. Note that a large number of SNPs are highly associated with both axes, reflecting strong population structure.

**Figure 18.**
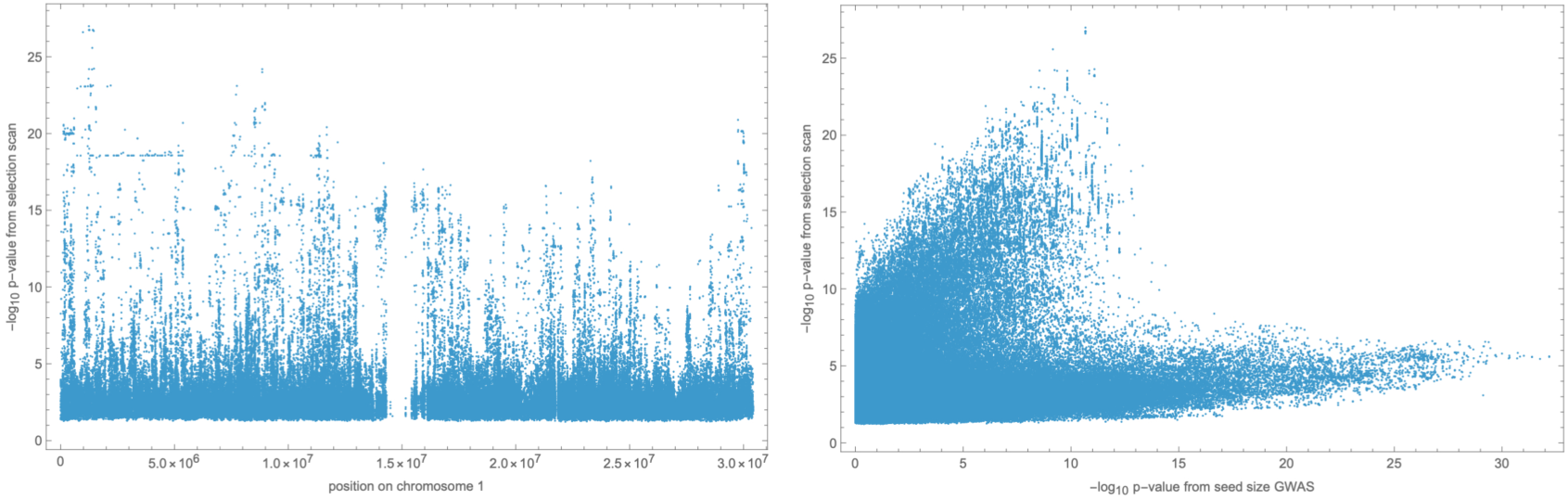
Left: results of selection scan for chromosome 1. Right: scatter plot of p-values from selection scan (all chromosomes) vs. seed size GWAS. Figure 18—figure supplement 1. Zoom-in of selection scan for chromosome 1 peak. Figure 18—figure supplement 2. Result of selection scan for chromosomes 2–4. Figure 18—source data 1. Results of selection scan.

However, as noted above, high fitness is not perfectly correlated with population structure, and while our SNP matrix contains 63 (out of 1,240,486) SNPs in multiple regions on all chromosomes distinguishing the B accessions from all other accessions, there are only 9 SNPs in a single 500 kb region on chromosome 1 that distinguish the B accessions plus the 3 other high-fitness beach accessions mentioned above from all other accessions. This region underlies the highest peak in the selection scan (***Figure 18***), and is centered on two plausible candidate genes: *YUCCA3* (AT1G04610), associated with auxin biosynthesis and early root growth (***Zhao et al., 2001***); and *MEE4* (Maternal Effect Embryo Arrest 4, AT1G04630), a mitochondrial protein with several embryo- and seed-related phenotypes (***Pagnussat et al., 2005***; ***Dong et al., 2016***)

## Discussion

This study was designed to look for evidence of local adaptation in Swedish *A. thaliana*, and to investigate whether selection experiments would reveal important factors that had been missed by traditional common-garden experiments. It was successful in both respects. Consider:

- A subset of southern S1 accessions experienced severe overwinter mortality (and decreased fecundity among survivors) in both northern sites in one year of our common-garden experiments, presumably reflecting a harsh winter (***Figure 3***).
- Slugs attacked one southern site in one year of our common-garden experiments, almost exclusively feeding on S1 and S2 plants, dramatically increasing overwinter mortality and decreasing fecundity among survivors (***Figure 4***).
- Accessions tended to have higher fecundity closer to home. Thus S1, S2, and B accessions had higher fecundity in southern than in northern sites, and the reverse was true for N accessions (***Figure 10***).
- This notwithstanding, S1 and S2 accessions generally had higher fecundity than N accessions across sites, and B accessions universally had low fecundity (***Figure 9***).
- Regardless of location, B accessions had a massive fitness advantage in all selection experiments, presumably related to seedling establishment (***Figure 15***).

The implied strength of selection was very large: in both the overwinter survival (***Figure 3***) and the selection experiment data (***Figure 15***), over five-fold differences in fitness between accessions were common.

We interpret these seemingly contradictory observations as reflecting at least three different evolutionary trade-offs: resistance (to stress) vs. high fecundity; seed quality (good seedling establishment) vs. seed quantity (high fecundity) and, finally; north vs. south—for unknown reasons, accessions tend to have higher fecundity closer to home. The latter sounds like textbook local adaptation, but this is not the case because even though N accessions have relatively higher fecundity in the north than in the south, they still have lower fecundity than S1 and S2 accessions. Without other factors influencing fitness, they would be outcompeted.

It is of course easy to see how a trade-off between resistance and fecundity could maintain variation among populations. Although we only have data for two years, our results for S1 and S2 vs. N are reminiscent of those of ***Oakley et al.*** (***2023***), who, through years of common-garden experiments using a RIL population generated by crossing an Italian and Swedish accession (from the same region as our N accessions), have found a similar trade-off under which Italian genotypes generally have an advantage due to higher fecundity, but suffer much higher mortality in the north in some years. In our experiments we see that while S1 and S2 accessions generally have higher fecundity, they are more sensitive to both slug and (presumably) weather-induced damage, resulting in increased mortality as well as decreased fecundity among survivors. However, to argue that this trade-off *actually* maintains variation would require much more information about the intensity of weather- and herbivory-induced stress on the requisite spatial and temporal scales (both of which are unknown)—and we would also need to know how relevant these common-garden estimates of survival and fecundity are to total fitness.

The answer to the latter question is far from obvious, as the striking results from our selection experiments make clear. Totally unexpectedly, we found that B accessions from the very south of Sweden had a fitness advantage in all sites, which is especially notable given their consistently low fecundity in our common-garden experiments. The implied selection strengths are enormous, with accessions increasing their frequency up to 20-fold over little more than a single life-cycle. While we do not know what caused these fitness differences, it is reasonable to hypothesize that it involves components not assayed in the common-garden experiments, in particular seedling establishment and competition (***Postma and Ågren, 2016***). This hypothesis is supported by the observation that fitness is associated with seed size, a trait that is highly likely to be important for seedling establishment, and is also largely genetically determined (***Figure 15***).

A trade-off between seed size and seed number is not surprising, but what could maintain variation for such traits? We would suggest that classical *r*/*k* selection plays a role (as has also been suggested by ***Bastias et al., 2024***; ***Lo et al., 2024***). The S1 and S2 accessions were mostly collected in small, ephemeral patches directly or indirectly created by human activity—a highly uncertain environment that might be expected to favor high fecundity and hence small seeds (*r*-selection). The N accessions, on the other hand, were mostly collected in patches created by ongoing natural erosion on south-facing slopes, many of which had harbored *A. thaliana* for at least a century according to local floras, and the B accessions, finally, were collected on extensive Baltic beaches—ancient, naturally disturbed habitats, maintained by the sea. Both environments might be expected to favor higher-quality seed (*k*-selection), perhaps to an extreme extent in the predictably harsh beach environment. Under this model, the B accessions would out-compete other accessions in almost any patch, only to go extinct when the patch disappears (*e.g.*, due to being overgrown by grass, or run over by a plow). The optimal strategy is determined by the frequency of disturbance.

To summarize, while we uncovered plenty of evidence of fitness-related variation consistent with local adaptation, we were also reminded of how complex the adaptive landscape is, and how difficult it is to measure fitness. Different experiments measure different fitness components and hence often produce contradictory results. Even our selection experiments, which included the full life cycle, measured total fitness conditional on being in a plot where competition is possible. If the kind of *r*-selection discussed above is important, then dispersal ability and seed banks would also play a major role (see, *e.g.*, ***Fakheran et al., 2010***). Designing experiments that include these fitness components would be challenging even if we knew the relevant spatial and temporal scales, which we assuredly do not (for other examples, see ***de la Mata et al., 2024***; ***Schmitz et al., 2024***). Certainly, a deep understanding of the natural history and ecology of the study organism is essential.

Our study also illustrates the challenges facing efforts to understand the "genetic architecture of adaptation" or, even more ambitiously, evolutionary change from the molecular to the ecological level (***Rockman, 2012***; ***Travisano and Shaw, 2013***; ***Hoban et al., 2016***). Leaving aside, for the moment, the fact that we do not know how to measure fitness, and focusing instead on fitness-related traits (which can surely be measured, as we have shown in this study), this program relies, explicitly or implicitly, on genetically dissecting traits in order to quantify the relative contribution of genetic polymorphisms, statistically or, more ambitiously, down to molecular causality. The latter is of course only possible for the subset of traits where sufficiently large-effect polymorphisms exist—a category that clearly excludes many complex traits (***Rockman, 2012***; ***Boyle et al., 2017***).

Unfortunately, dissecting adaptive traits is very challenging even when large effects do exist. Consider our results for slug herbivory, which demonstrated negative effects of fall herbivory on over-winter survival (***Figure 4***), as well as fecundity among the survivors in spring (***Figure 9***—***figure Supplement 1***). Glucosinolates are known to deter herbivores, and glucosinolate profiles in *A. thaliana* are largely controlled by two major polymorphisms, at the *AOP* and *MAM* gene clusters, only the former of which is segregating at high frequency in Sweden (***Kliebenstein et al., 2001***; ***Katz et al., 2021***; ***Gloss et al., 2022***). We confirmed that these profiles are strongly associated with both slug damage and over-winter survival (***Figure 6***), and we also noted that one of the top GWAS hits for over-winter survival contains the *AOP* cluster (***Figure 5***—***figure Supplement 3***). The chain of causation is thus reasonably clear: a major polymorphism at *AOP* affects glucosinolate profiles, which affect slug damage, which affects over-winter survival and fecundity, both of which are surely fitness-related traits. However:

- While the *AOP* association is indeed one of the strongest associations for over-winter survival, it is one of several, and it is far from genome-wide significant after correcting for population structure. Without the benefit of prior information, we would not have identified it. And it was not strongly associated with fecundity.
- While the existence of a causal relationship between the *AOP* polymorphism and over-winter survival and fecundity seems clear, genome-wide linkage disequilibrium makes unbiased estimates of its marginal contribution very difficult to obtain.
- As discussed above, it is very unclear how fitness estimates from our experiments relate to overall adaptation in Sweden. Specifically, there is reason to believe that the kind of slug damage seen in one year in one field site might be more common in a relatively densely planted common garden experiment than in nature.

There are no easy fixes to any of these problems. With respect to the first two points, genome-wide linkage disequilibrium is a direct consequence of population structure (as first realized by ***Wahlund, 1928***), and leads to biased effect-size estimates across the genome due to correlations with causative polymorphisms. Selfing further aggravates the problem (***Nordborg, 2000***), but it is important to note that extensive linkage disequilibrium *per se*, whether due to selfing or population structure, is not the main problem. The really serious problem is that local adaptation will, by definition, strengthen correlations between causative loci and population structure—leading to stronger genome-wide confounding, as well as increased linkage disequilibrium between the causative loci themselves (reflecting selection as well as population structure, see ***Ohta, 1982***), making unbiased marginal effect-size estimates difficult to obtain. Simple genetic traits, like anthocyanin production (***Figure 11***), are not strongly affected because there is not much of a genetic background to confound estimates.

The problem is well known to plant breeders, and many statistical solutions have been proposed (reviewed in ***Vilhjálmsson and Nordborg, 2012***). However, there are limits to what statistics can achieve, as illustrated here. The alternative approach is of course genetic crosses, which break up long-range associations very effectively, but typically lack resolution to identify causal polymorphisms. They also cannot be used directly to estimate genetic architecture, because it is a property of the population in which it is measured. For example, as mentioned above, a loss-of-function allele of *CBF2* plays a major role in freezing tolerance in a mapping population generated by crossing a Swedish and an Italian accession (***Ågren et al., 2013***; ***Gehan et al., 2015***; ***Lee et al., 2024***), but probably no role in natural European populations as the allele is very rare.

Finally, we can dispense with mapping altogether and look directly for signs of selection using a variety of methods from population genetics. However, it is not clear how one quantifies genetic architecture based on such results.

Regardless of all this, and to end on a positive note, what our study demonstrates is the value of field studies and the importance of phenotypes. Before trying to understand adaption on the genetic (or molecular) level, we must understand it on the phenotypic level.

## Methods

### Genotypes

All 200 accessions were re-sequenced to confirm identity. This uncovered minor discrepancies, most likely reflecting residual heterozygosity in the the original lines, and we generated a new SNP matrix to reflect this (and improvements in SNP-calling algorithms, see ***Brachi et al., 2022***).

In addition, during the course of writing this paper, we discovered that two of our accessions are not what they are supposed to be. These two accessions (1435 and 6180) are annotated as originating from northern Sweden, but several lines of evidence demonstrate that this is false:

1. their SNP genotypes do not match early genotypes for these accessions ***Horton et al.*** (***2012***), nor genotypes in old crosses made with these accessions;
2. in our analysis of population structure, they bear no resemblance to other N accessions and instead cluster with S1 accessions;
3. in particular, the pattern of haplotype sharing indicates that they are very closely related (possibly part of an inbred sibship) with accession 5829, supposedly collected at Ales Stenar in the extreme south of Sweden.

Since all of our analyses are based on the actual genotypes, our results are unaffected by this.

### Population structure

To explore patterns of population structure in our sample of 200 accessions from Sweden, we generated a set of 124,071 LD-pruned SNPs with MAF ≥ 3%. MAF-filtering and LD-pruning were performed in plink v1.9 (***Chang et al., 2015***) using the options -maf set to 0.03, and –indeppairphase with a window of 5 kb, a step size of 10 SNPs and a *r*^2^ threshold of 0.2.

Genetic distances were calculated from these SNPs using the read_plink function from the R package genio v1.1.2 to read in the data, and the dist.gene function from the package ape v5.7-1 (***Paradis and Schliep, 2019***). The neighbor-joining tree was created with the nj function. Admixture analyses used the R package LEA 3.12.2 (***Frichot and Francois, 2015***). Specifically we used the snmf function and performed 10 runs for *k* ranging from 1 to 8. Ploidy was set to 2, and the snmf regularization parameter alpha to 100. For each *k* we picked the run with the minimum cross entropy.

To present the results from our study we picked *k* = 4 as it the largest value for which “pure" accessions are found (***Figure 1***—***figure Supplement 1***). Accessions were assigned to one of four group based on their largest components. As shown in ***Figure 19***, many accession are intermediate between S1 and the other three groups, and there is also some admixture between S1, B, and N.

**Figure 19.**
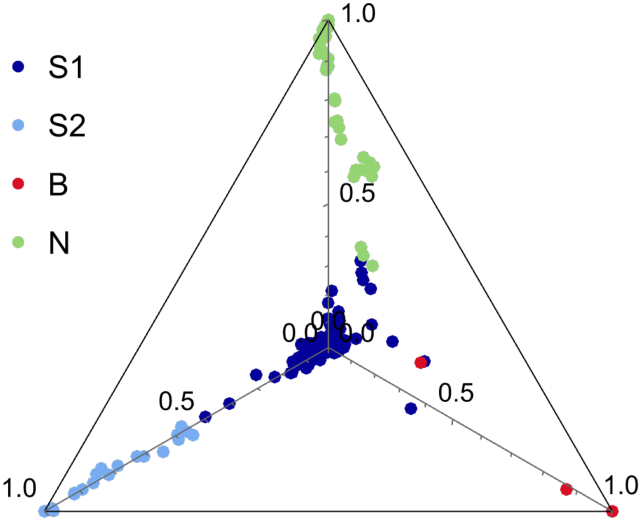
Accessions plotted in the simplex space generated by the admixture proportions.

### Common-garden experiments

#### Experimental design

These experiments are explained in detail in ***Brachi et al.*** (***2022***). Briefly, seed for 200 accessions of *A. thaliana* from Sweden were produced together in a randomized design in the greenhouse of the University of Chicago under long-day conditions with a 12-week vernalization period at 4°C to induce flowering. Pots were continually randomized until flowering to avoid position effects. The resulting seeds were then used to set up the experiments, repeated over two seasons in four locations in Sweden:

NA South-facing slope near Ådal (lat. 62.862, lon. 18.331)

NM Agricultural field near Ramsta (lat. 62.85, lon. 18.193)

SR Agricultural field near Rathckegården (lat. 55.906, lon. 14.260)

SU Agricultural field near Ullstorp (lat. 56.067, lon. 13.945)

Each experiment was organized in three complete blocks, each including eight replicates of each accession. Accessions replicates were randomized within blocks. Seeds were sown in trays of 66 pots, each measuring 4 cm in diameter, in a mix of 90% standard greenhouse soil and 10% local soil. Seedlings were allowed to establish outside under shelter and were thinned to single plant per pot before being moved to the field sites and laid on tilled soil. Plants were watered once upon installation. The same dates were used in both years (***Table 1***).

**Table 1.** Sowing and installation date for common-garden experiments. Blocks A–C were sown with two-day intervals.

|  | 2011 |  | 2012 |  |
| --- | --- | --- | --- | --- |
|  | sowing (blocks A–C) | field installation | sowing (blocks A–C) | field installation |
| NA | 8/8–8/12 | 8/25 | 8/8–8/12 | 8/25 |
| NM | 8/7–8/11 | 8/24 | 8/7–8/11 | 8/24 |
| SR | 9/1–9/5 | 9/18 | 9/1–9/5 | 9/18 |
| SU | 8/31–9/4 | 9/17 | 8/31–9/4 | 9/17 |

#### Phenotyping

We scored the presence and absence of each plant in fall and in spring, just after snow melt, and used these data to compute overwinter survival as a simple binary variable. Note that in the fall of 2012 we collected from 0–2 replicates per block to study gene expression and methylation (data not used in this study), and in the spring of 2012 and 2013, we also sacrificed two replicates (per accession and block) for DNA extraction and microbial community analyses (***Brachi et al., 2022***). The data included with this paper reflect this.

In one experiment (SR 2011-12), the plants suffered damage from slugs, which we scored in November 2011, using *ad hoc* scale: 0 (no plant), 1 (undamaged plant), 2 (mild damage) and 3 (extensive damage).

In addition, we photographed trays in both fall and spring. The images taken in the fall were used to investigate rosette sizes and purpleness using high-throughput image analyses (see the Rosette purpleness section, below).

The remaining replicates for each accessions were left to complete their life cycle in the field. With very few exceptions, all plants flowered within a two-week period in spring in each experiment (flowering is several weeks later in the north), and our records were insufficiently fine-grained to consider genetic difference.

When seeds were mature in late spring, all plants remaining in the experiments were harvested for fecundity estimation. This was achieved using high-throughput image analysis, validated by hand measurements (details in ***Brachi et al., 2022***).

### Analysis

#### Survival

The overwinter survival data were analyzed in a generalized mixed model framework using the lme4 R package v 1.1-35.5 (***Bates et al., 2015***). Specifically we used the glmer function with optimizer set to bobyqa and the iterations to 2 × 10^5^ with option maxfun to fit models of increasing complexity. The simplest model included a single random block effect to explain survival, and the most complex model is described by:

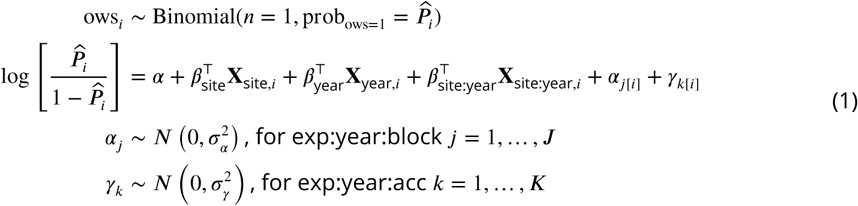

In this model, ows_i_ describes the overwinter survival of the i^th^plant: 0 for dead plants, 1 for plants still alive. ***P****^_i_* is the probability of surviving winter and is modeled using a logit link function. *α* is the model intercept and βs are vectors of fixed effect coefficients for sites, years, and their interaction. Block effects within sites and years are captured by a random intercept *α*_j_, which is assumed to follow a normal distribution. Accession effects within sites and years are captured by a second random intercept term ⊲_0_, also assumed to follow a normal distribution. This model fitted the data better (based on AIC, BIC and log-Likelihood) than a model without the random accession effect within site and year (⊲_0_). Accessions effects or BLUPS (Best Linear Unbiased Predictions, ⊲_0_) were computed from this model and used in GWAS.

To compare survival in the two northern sites in 2011-22 (***Figure 3***), we used a simple liability-threshold model, which also underlies the logit-transformation above. Briefly, we assume liability is Gaussian with accession i having a genetically determined liability N(µ_i_, σ_NA_) or N(µ_i_, σ_NM_) and that the probability of survival is the cumulative probability up to exposure τ_NA_ *>* τ_NM_ (reflecting the harsher environment in NM). Using this model, the relationship between the survival probabilities in NM and NA can be written

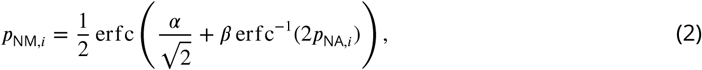

where *α* = (τ_NA_ – τ_NM_)/σ_NM_, *β* = σ_NA_/σ_NM_, and erfc is the complementary error function. We fit this equation to the data using NonlinearModelFit in Mathematica.

#### Herbivory damage

Slug herbivory (observed in SR 2011-12 only) was scored as "heavy", "moderate", and "no" damage. The analysis simply uses the proportion of plants per accession with heavy damage.

#### Rosette purpleness

Trays in the common-garden experiments were photographed as follows:

- SR on the 09/29/2011, 11/20/2012 and 11/25/2012
- SU on the 11/01/2011, 09/11/2011 and 11/22/2012
- NA and NM on 09/29/2012

Tray images were cropped by hand to the edge of the trays, and plants were then segmented automatically using custom scripts based on the EBImage R package (***Pau et al., 2010***). Briefly, the image segmentation starts by normalizing the images. Pixels are then clustered with the R function clara from the package cluster, with *k* = 3. The cluster with the highest average green value is considered to represent mostly plants. Pixels from the other clusters were masked and the segmentation of plant was refined by successive erosion, expansion and blurring. The result consists of tray images with up to 66 plants on a white background. Images were all manually checked and the segmentation corrected when needed in Photoshop CS6 (*i.e.*, separating touching plants or removing remains of the background). These images were then used to measure the color composition of individual rosettes (and their size). First, white and black pixels were removed from the segmented images as they carry no information. Colors from the whole image were then reduced to 16 color groups, using k-means clustering (based on the RGB values). The frequency spectrum of 16 colors was then computed for each plant on the image and each plant was assigned to a position on the tray in terms of rows (1-6) and columns (1-11). The resulting table consists, for each tray, of the plant coordinates on the tray (used to match with accession identity) and 16 columns of pixel counts corresponding to the 16 colors. Simultaneously, the RGB values of the 16 colors identified on each tray were recorded in another table.

The RGB values of all observed colors across all trays in all experiments (16 colors per tray, amounting to 17,152 unique colors) were then summarized using a principal components analysis of the normalized R, G and B values. In this analysis the purple to green gradient is well captured by the first component, which explains 75% of the variance. The coordinates of the 16 colors from each tray along this component are then multiplied by the proportions of pixel of these colors found in each plant. The resulting vector quantifies the purpleness in 32,154 rosette images.

To investigate the effects of genetic variation on plant purpleness, we modeled purpleness using the R package lme4, using the following model:

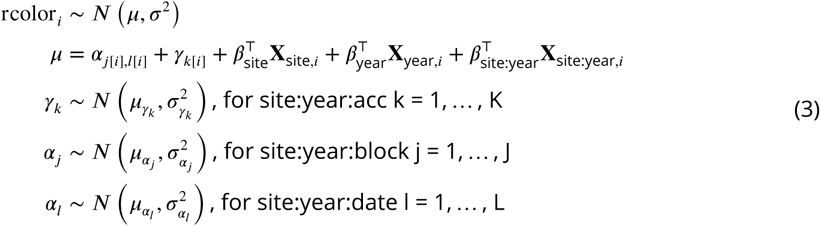

In this model, rcolor_i_ is the purpleness of the i^th^plant. The terms of the model are largely identical to the models used above. Purpleness variation is explained by fixed effects for site and year effects and their interactions (*β*), as well as random intercept effects for accessions (⊲_0_) and blocks within sites and years (*α*_j_). We added a random intercept terms for photography date (*α*_l_) as some experiments were photographed twice the same year. This model was used to generate BLUPs used in GWAS and to investigate genetic correlations with other traits.

#### Fecundity

The per-plant fecundity estimates were derived from image analysis of mature stems. Note that the raw areas occupied by mature plants on images (our fecundity estimate) was log-transformed (log_10_) prior to analyses. All plants for which we retrieved no stem, or only fragments were removed from this analysis (there are no zeros in the data).

Estimates were again modeled using a linear mixed model. We proceeded exactly as we did for the overwinter survival, starting from a simple model including only a random block effect, to incrementally complex models. Again, the most complex model provided a better fit (based on AIC, BIC and log-likelihood):

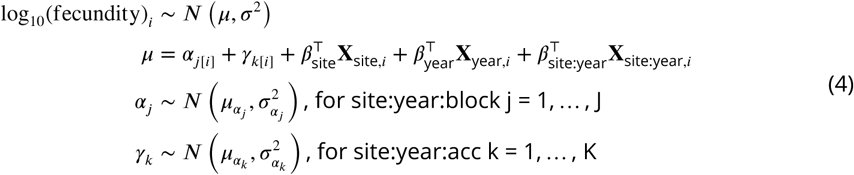

Terms in this model are described for other models. Model residuals were checked for heteroskedasticity and large deviation from normality.

As for overwinter survival, BLUPs were extracted for accessions within sites and years and used as phenotypes in GWAS. We also investigated patterns of variation among accessions across the eight combinations of sites and years using a principal component analyses.

#### GWAS

Accession effects (BLUPs) obtained from generalized linear models described above were used in GWAS. The SNP matrix used was filtered to include only bi-allelic SNPs with MAF ≥ 3%. Traits were mapped for sites and years separately using the generalized linear models (GLM, no control for population structure) and mixed-models (MLM, controlling for population structure using a kinship matrix) implemented in rMVP v1.0.6 (***Yin et al., 2021***). Custom plotting functions based on ggplot2 were used for generating Manhattan plots, including zoom-ins.

### Experimental evolution experiments

#### Installation

Our experimental evolution experiments were carried out at four sites, two in the north, and two in the south (***Figure 1***):

NA South-facing slope near Ådal (lat. 62.862, lon. 18.338)

NB South-facing slope near Barsta (lat. 62.87, lon. 18.381)

SR Agricultural field near Rathckegården (lat. 55.906, lon. 14.260)

ST Sandy beach/pasture near Tjörnedala (lat. 55.60, lon. 14.304)

Sites NA and SR were also used for common-garden experiments. The sites were chosen to be plausible *A. thaliana* habitats, but without *A. thaliana* visibly present. No vegetation surveys were made, but photos in ***Figure 1***—***figure Supplement 4*** and ***Figure 1***—***figure Supplement 5*** provide an idea of what the sites were like. No clearing of existing vegetation was done except in the SR site, a sandy agricultural field, which was initially cultivated, then left alone.

Within each site we delimited four to five replicate 1 m^2^ plots marked with large metal nails hammered into the ground until invisible. Thus the plots themselves were invisible, but could be relocated using a metal detector. Each plot consisted of 3 to 4 contiguous 60 × 40 cm subplots arranged to match the terrain. Using seeds from the same lots used for the common-garden experiments (see above), we prepared tubes containing 40 seeds for each of the 200 accessions using an Elmor C3 seed counter. Prior to dispersal, seed from each tube were mixed with sterile sand of about the same granularity as Arabidopsis seeds, and transferred to ϱ 20 ml plastic vials with holes drilled in the caps. This allowed for homogeneous dispersal of 40 × 200 = 8,000 seeds in each subplot, guided by a 60 × 40 cm frame with a string mesh delimitating 10 × 10 cm squares. The dispersal was repeated 3 times over three weeks in 2011 to maximize the chances of successful establishment. In the north (NA and NB), we dispersed seeds August 4th–6th, 15th–16th and 23rd–24th. In the south (SR and ST), we dispersed seeds August 28–30, September 7–8 and September 15–16. In total we thus dispersed 120 seeds per accession per m^2^.

#### Genotyping

Individual plants we sampled randomly using a grid in spring 2013. At least 70 plants were sampled per (surviving) plot in each plot (***Figure 1***—***figure Supplement 4*** and ***Figure 1***—***figure Supplement 5***). Multiplexed DNA libraries were prepared using the Illumina Nextera™ Kit. The standard Nextera library construction protocoal was changed to handle reduced volumes. Tagmentation reaction was set up to 2.5 µl final volume with 2.5 ng of input DNA. For PCR amplification and multiplexing we used Illumina Nextera Dual primers. Size selection and PCR cleanup were performed with Agencourt AMPure XP Beads (Beckman Coulter). After PCR enrichment, libraries were validated with Fragment Analyzer™ Automated CE System (Advanced Analytical) and pooled in equimolar concentration for 96X-multiplex. Libraries were sequenced on Illumina HiSeq™ V4 Analyzers using manufacturer’s standard cluster generation and sequencing protocols in 125 bp PE mode at the Vienna Biocenter Core Facilities using standard Illumina paired-end protocols. We produced on the order of 1-2 million reads for each sample with an average read-length of 100 bp. These reads were mapped to the reference TAIR10 genome using bwamem with default parameters. We genotyped 2.3 million previously identified SNP using bcftools with default parameters (base quality threshold of 30 and read mapping quality of 10). The nextflow pipeline for SNP calling is available at https://github.com/Gregor-Mendel-Institute/nf-haplocaller.

In order to assign each sample to one of the 200 experimental accession, the workflow summarized in ***Figure 20*** was used. We outline each step below.

1. Many samples were very small (collection was done using tweezers), and there was a clear risk of sampling the wrong species. To quickly eliminate such samples, we mapped reads to *A. thaliana* centromeric repeats and organellar genomes.
2. We estimated the minimum number of SNPs required for genotyping the samples using the SNPmatch “simulation” function (***Pisupati et al., 2017***). With the exception of two pairs of accessions, 5k SNPs random were sufficient to distinguish all accession. To be conservative, we set a threshold of 10k SNPs.
3. We used SNPmatch and an empirically derived mismatch threshold of 0.015 (see ***Pisupati et al., 2017***) to assign samples to experimental lines.
4. Of the roughly 1/4 of samples that did not match an experimental accession, roughly 1/3 showed extensive heterozygosity that could either reflect recent outcrossing or sample contamination (the quality of our data was not sufficient to investigate this further, nor were we able to identify parents of putatively outcrossed individuals). The remaining 2/3 were inbred accessions that we classified as “natives".
5. Of the 3/4 of samples that did match an experimental accession, 19% matched an accession that might also have been present as native in the experimental site where the sample was taken (a “potential native"; see next section) while the remaining 81% were unambiguously experimental samples.

#### Potential natives

The observation that all four experimental sites contained obvious natives alerted us to the possibility that our results could be biased by the presence of cryptic natives, *i.e.*, native members of one of our experimental accessions. This would be far more likely to happen for an experimental accession that was originally collected close to the experimental site, because, with the notable exception of North America, where the species is recently introduced, *A. thaliana* is characterized by strong isolation-by-distance and the probability of finding identical individuals decays rapidly with distance (***Platt et al., 2010a***).

This pattern can be seen in ***Figure 21***, where we plot isolation-by-distance for the old data by ***Platt et al.*** (***2010a***), and also for the new “beach"-centered collection used to examine the distribution of seed size (***Figure 17***). The new data confirm the conclusion of ***Platt et al.*** (***2010a***), but has higher resolution due to much denser sampling. Identical genotypes are very rarely seen more than a few km apart, although exceptions exist in the south (where they are likely due to human dispersal). In the almost continuous beach population we sampled, the probability effectively declines to zero (*p* = 3.9 × 10^–6^) at 5.6 km.

**Figure 20.**
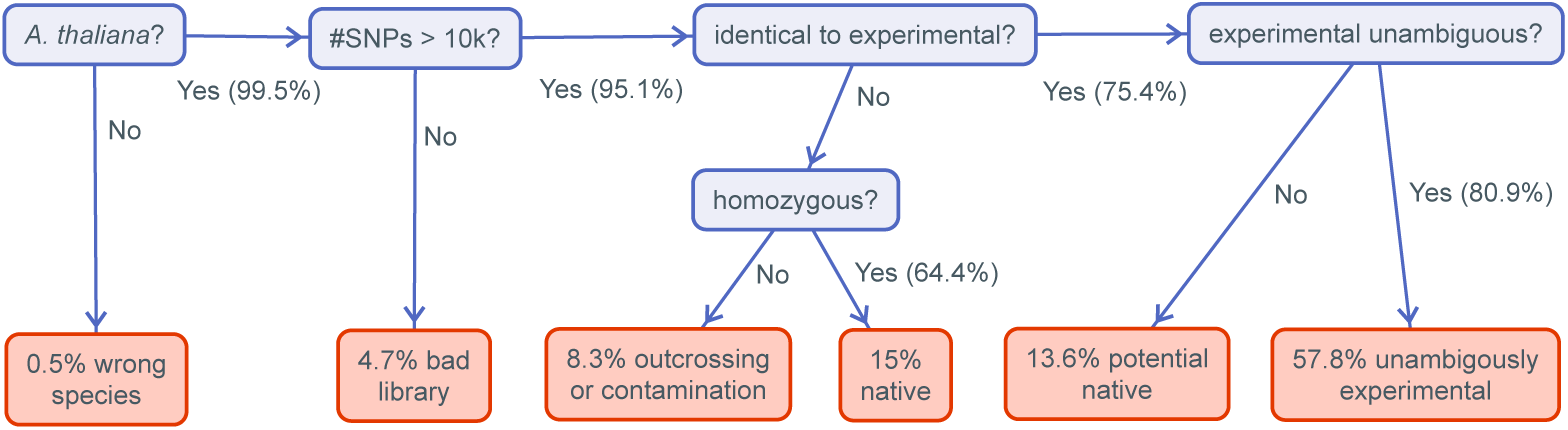
Workflow for classifying selection experiment samples. Figure 20—figure supplement 1. The number of samples in each category per site and plot.

**Figure 21.**
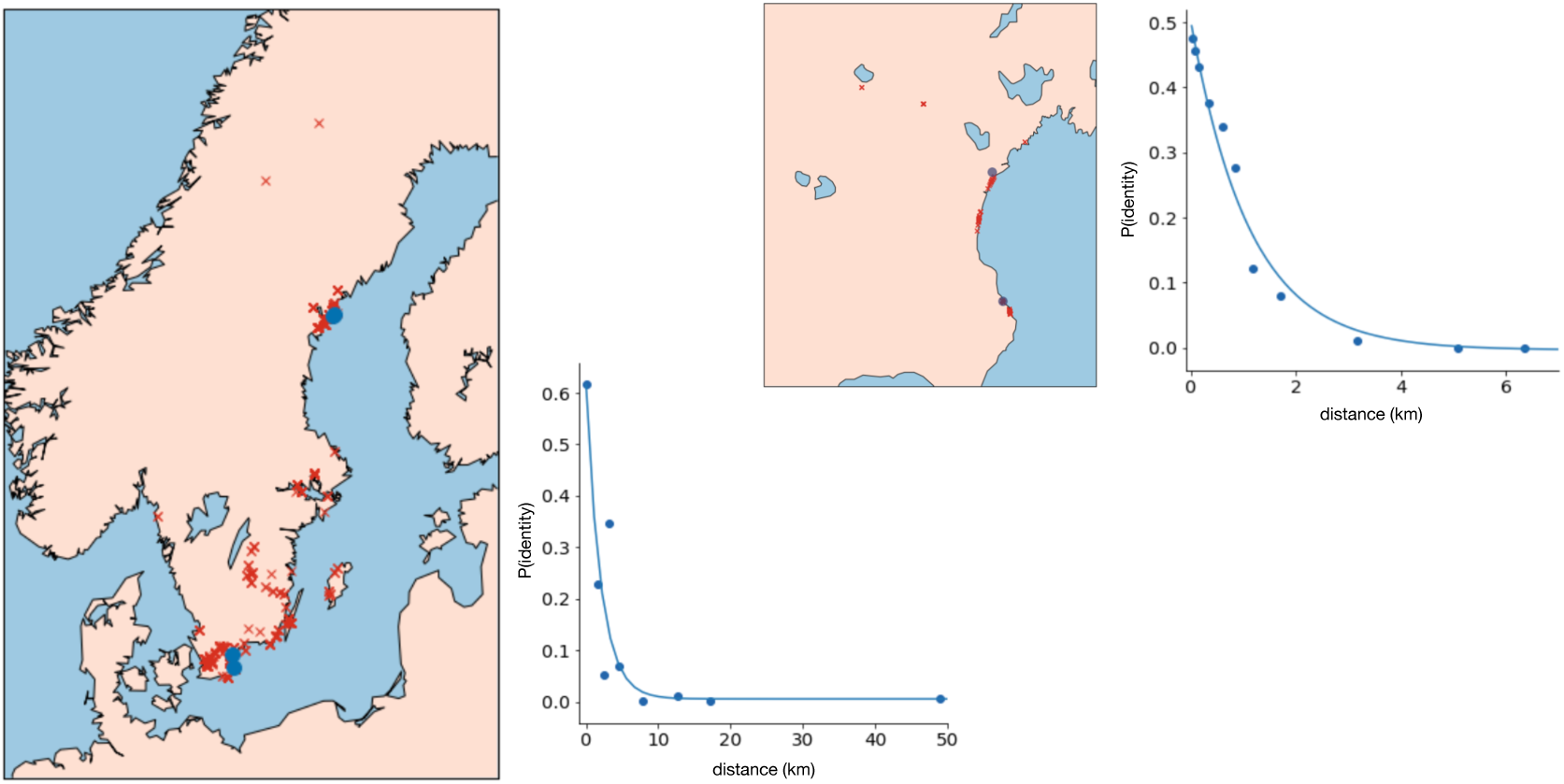
Isolation-by-distance in Swedish *A. thaliana*. On the left, sample locations and the decay in pairwise identity as a function of distance using the data of ***Platt et al.*** (***2010a***); on the right, the same plots for the new samples from Skåne (Figure 17). The blue dots show the sites used in the evolution experiments.

Taking 5.6 km as a cutoff, we see that results for a total of 11 accessions could have been confounded by cryptic natives in the northern sites, and results for 8 accessions could have been confounded at the ST site. No experimental accession was sampled close enough to the SR site for it to be plausible that result were confounded by cryptic natives.

As shown in ***Figure 13***, we find convincing evidence that two data points (the Bar1 accession in NB, and the Vår2-6 accession in ST) were indeed confounded. However, more importantly, the main result of these experiments, namely that the same beach accessions outperformed inland accessions in all sites, could not possibly be explained by this kind of confounding.

#### Seed size

Seed size measurements (taken as described in ***Clauw et al., 2022***) were available for most of the accessions used in this study. ***Figure 15*** uses the published measurements, which were taken on seeds produced in the lab, but it is important to note that these measurement are highly correlated with those produced in the field, and also with estimates from other labs, in completely independent experiments (***Figure 15***—***figure Supplement 1***). This demonstrates that the seed size variation among accessions observed here is largely genetic, and strongly suggests that large seed size is a characteristic of the beach accessions.

Further evidence of this is provided by the observation that seed size in a new sample of 246 beach accession mirrored what we had already seen. Seed size for these accessions was measured using Boxeed (Labdeers, Czech Republic).

#### Modeling allele-frequency change

The experimental sites were sown in fall 2011, and seedlings were allowed to establish normally and set seed in spring/summer 2012, giving rise to a second generation that germinated in fall 2012. Survivors were then sampled in spring 2013. While we cannot rule out that some samples might be dormant survivors of the original sowing rather than descendant of this generation, we believe their contribution would be minor, as there was no evidence of dormancy in the originally sown seeds, and a substantial flowering population in 2012 in all sites.

Either way, from the perspective of understanding how allele-frequencies would change in the absence of selection, the population of (experimental) individuals in spring 2013 can be modeled as Wright-Fisher multinomial sample of size N_9_from a starting population where all 200 accessions had equal frequency. At the level of SNPs, the population frequency of an allele with starting frequency *p*_0_ will thus be binomially distributed with parameters N_e_ and *p*_0_. This frequency is what we then estimate using *bona fide* binomial sampling with (known) sample size *n*. Let Δ*p* be the difference between our estimate and the original allele frequency *p*_0_. In the absence of selection, using basic probability theory, we have E[ςx] = 0 and

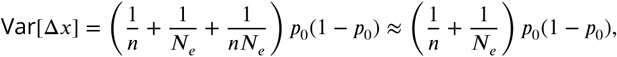

*i.e.*, allele frequencies should only change due to random sampling (in this case “drift" plus literal random sampling). As shown in ***Figure 22***, mean Δ*p* is indeed close to zero, and the variance is a linear function of *p*_0_(1 –*p*_0_) as predicted by the equation above. Since we know the sample size *n*—it was 171, 199, 208 and 137 for NA, NB, SR, and ST, respectively—we can estimate the corresponding N_e_to be 17, 70, 34, and 16 from the fitted slope in the figure.

**Figure 22.**
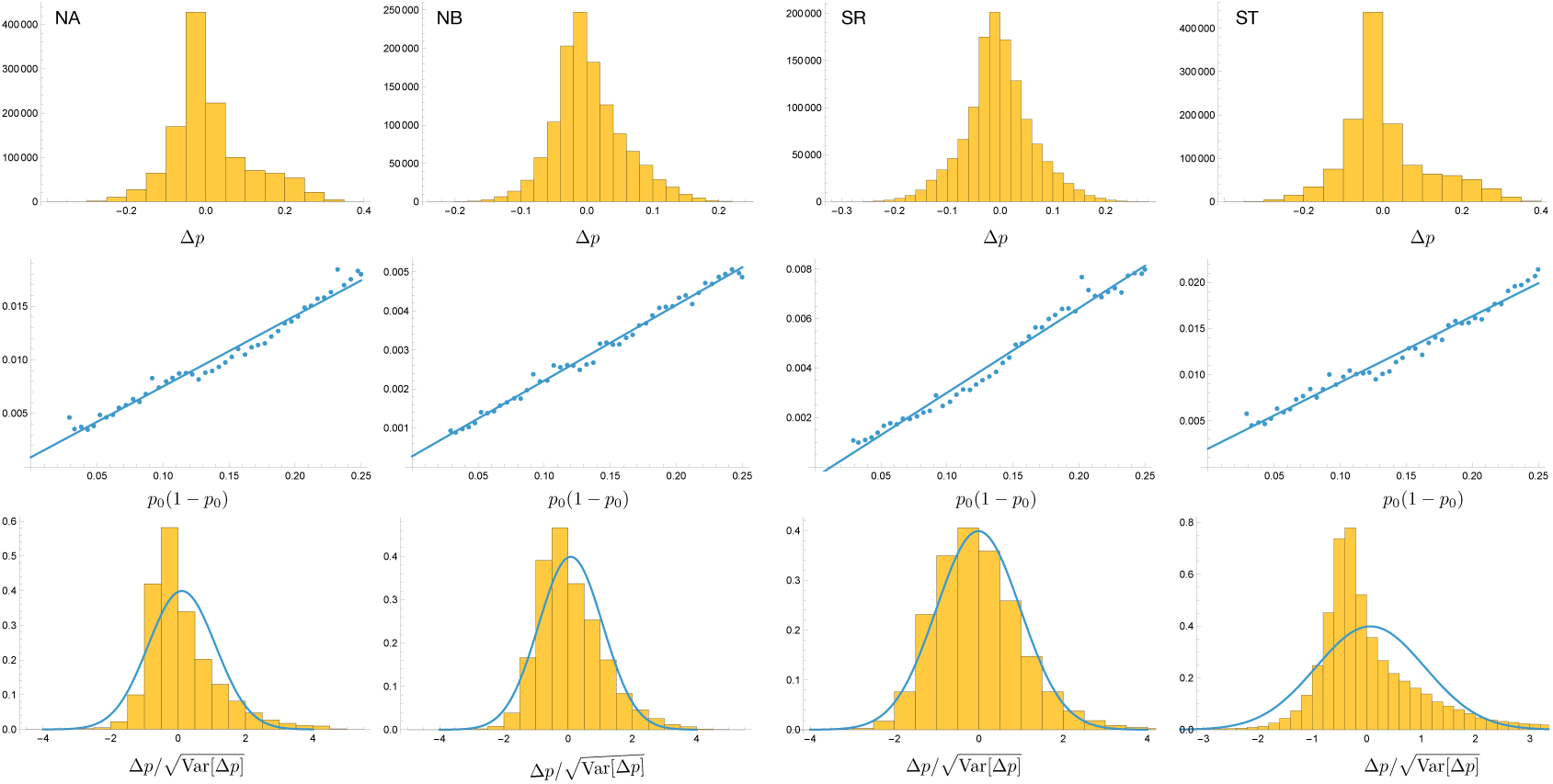
Top: the distribution of ς*p* in the four selection experiments. Middle: the variance of ς*p* as a function of *p*_0_(1 – *p*_0_). Bottom: the distribution of ς*p* scaled by its standard deviation. The curves are PDFs of normal distributions with the observed mean and variance 1.

To identify Δ*p* values too large to be due to drift, we divide them by the estimated standard deviation for the corresponding Δ_0_, and calculate the significance of deviations from zero using a standard normal distribution. Note that the mean is not zero, reflecting the fact that allele-frequencies are biased towards a German reference genome. More importantly, there are clear genome-wide deviations reflecting strong selection and population structure (as is also evident in ***Figure 18***).

## Acknowledgments

M.N. dedicates this paper to the memory of Malte Jönsson (***Figure 23***), who helped collect most of the accessions used in this study and provided invaluable knowledge of the land and its use. Further thanks go to Mia Holm for her hospitality and wonderful dinners after hard work in the field as well as help during harvesting; to Einar Holm for helping with field work and taking photos of harvested plants; to Ingalill Thorsell and the Drakamöllan staff for hosting retreats, and, finally; to the Kleen family, the Öhman family, Nils Jönsson (***Figure 23***), and the Rathckegården farm for allowing us to install our experiments on their land.

**Figure 23.**
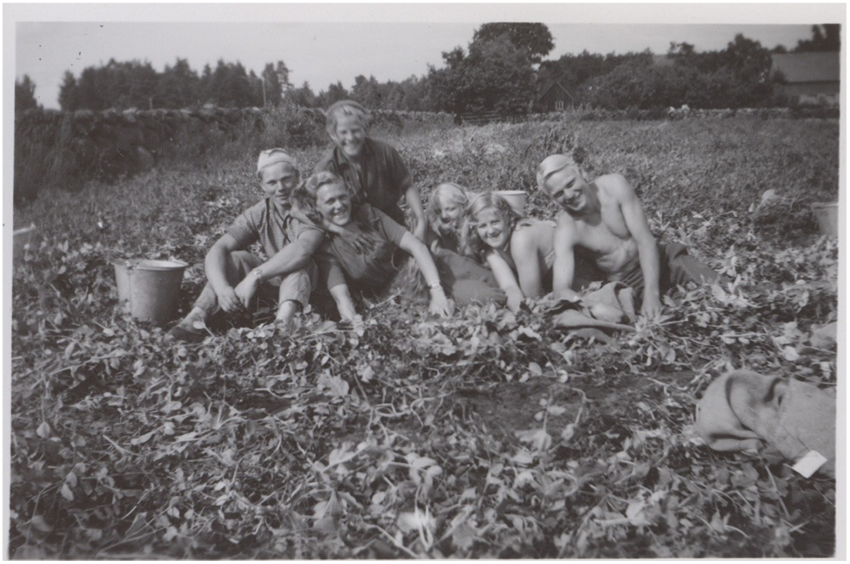
Brothers Nils (left; 1929–2025) and Malte (right; 1931–2022) Jönsson with friends carrying out field experiments with peas near the SU site shortly after WWII.

M.N. wrote much of this paper while on a sabbatical in Molly Przeworski’s lab at Columbia University, and thanks all members of her lab, as well as the labs of Guy Sella and Peter Andolfatto for their hospitality and feedback. Thanks also go to Man Yu from the C.D. lab, who helped generate stem images used for seed-set estimates and manual seed-set estimate, and to Tom Ellis and Jon Ågren for discussions and extensive comments on the manuscript.

This work was funded by a grant from the National Health Institute (Grant R01 GM 083068) to J.B., M.N., and C.D.; by an ERC AdvG (no. 268962—“MAXMAP") to M.N.; by a Dropkin Foundation Fellowship to B.B. J.B. was further supported by the University of Chicago and New York University, while M.N. was supported by the Gregor Mendel Institute and the Vienna BioCenter Core Facilities. B.B. received the support of the European Union in the framework of the Marie-Curie FP7 COFUND People Programme, through the award of an AgreenSkills/AgreenSkills+ fellowship (under Grant Agreement 267196).

## References

1. 1001 Genomes Consortium. 1,135 Genomes Reveal the Global Pattern of Polymorphism in *Arabidopsis thaliana*. Cell. 2016 Jul; 166(2):481–491.

2. Ågren J, Oakley CG, McKay JK, Lovell JT, Schemske DW. Genetic mapping of adaptation reveals fitness tradeoffs in *Arabidopsis thaliana*. Proceedings of the National Academy of Sciences. 2013; 110(52):21077–21082.

3. Ågren J, Schemske DW. Reciprocal transplants demonstrate strong adaptive differentiation of the model organism *Arabidopsis thaliana* in its native range. New Phytol. 2012 Jun; 194(4):1112–1122.

4. Alexander DH, Novembre J, Lange K. Fast model-based estimation of ancestry in unrelated individuals. Genome Res. 2009 Sep; 19(9):1655–1664.

5. Antonovics J, Clay K, Schmitt J. The measurement of small-scale environmental heterogeneity using clonal transplants of Anthoxanthum odoratum and Danthonia spicata. Oecologia. 1987 Mar; 71(4):601–607.

6. Atwell S, Huang YS, Vilhjálmsson BJ, Willems G, Horton M, Li Y, Meng D, Platt A, Tarone AM, Hu TT, Jiang R, Muliyati NW, Zhang X, Amer MA, Baxter I, Brachi B, Chory J, Dean C, Debieu M, de Meaux J, et al. Genome-wide association study of 107 phenotypes in *Arabidopsis thaliana* inbred lines. Nature. 2010 Jun; 465(7298):627–631.

7. Bastias CC, Estarague A, Vile D, Gaignon E, Lee CR, Exposito-Alonso M, Violle C, Vasseur F. Ecological trade-offs drive phenotypic and genetic differentiation of *Arabidopsis thaliana* in Europe. Nat Commun. 2024 Jun; 15(1):5185.

8. Bates D, Mächler M, Bolker B, Walker S. Fitting Linear Mixed-Effects Models Using lme4. Journal of Statistical Software. 2015; 67(1):1–48. doi: 10.18637/jss.v067.i01.

9. Boyle EA, Li YI, Pritchard JK. An Expanded View of Complex Traits: From Polygenic to Omnigenic. Cell. 2017 Jun; 169(7):1177–1186.

10. Brachi B, Filiault D, Whitehurst H, Darme P, Le Gars P, Le Mentec M, Morton TC, Kerdaffrec E, Rabanal F, Anastasio A, Box MS, Duncan S, Huang F, Leff R, Novikova P, Perisin M, Tsuchimatsu T, Woolley R, Dean C, Nordborg M, et al. Plant genetic effects on microbial hubs impact host fitness in repeated field trials. Proc Natl Acad Sci U S A. 2022 Jul; 119(30):e2201285119.

11. Chang CC, Chow CC, Tellier LCAM, Vattikuti S, Purcell SM, Lee JJ. Second-generation PLINK: rising to the challenge of larger and richer datasets. Gigascience. 2015 Dec; 4(1):7.

12. Clausen J, Keck DD, Hiesey WM. Experimental studies on the nature of species. I. Effect of varied environments on western North American plants, vol. 520 of Publication. Washington, DC: Carnegie Institute of Washington; 1940.

13. Clauw P, Kerdaffrec E, Gunis J, Reichardt I, Nizhynska V, Koemeda S, Jez J, Nordborg M. Locally adaptive temperature response of vegetative growth in *Arabidopsis thaliana*. bioRxiv. 2022 Feb; p. 2022.02.15.480488.

14. Dong CJ, Wu AM, Du SJ, Tang K, Wang Y, Liu JY. GhMCS1, the cotton orthologue of human GRIM-19, is a subunit of mitochondrial complex I and associated with cotton fibre growth. PLoS One. 2016 Sep; 11(9):e0162928.

15. Fakheran S, Paul-Victor C, Heichinger C, Schmid B, Grossniklaus U, Turnbull LA. Adaptation and extinction in experimentally fragmented landscapes. Proceedings of the National Academy of Sciences. 2010; 107(44):19120–19125.

16. Fournier-Level A, Korte §, Cooper MD, Nordborg M, Schmitt J, Wilczek AM. A map of local adaptation in *Arabidopsis thaliana*. Science. 2011; 334(6052):86–89.

17. Frachon L, Libourel C, Villoutreix R, Carrère S, Glorieux C, Huard-Chauveau C, Navascués M, Gay L, Vitalis R, Baron E, Amsellem L, Bouchez O, Vidal M, Le Corre V, Roby D, Bergelson J, Roux F. Intermediate degrees of synergistic pleiotropy drive adaptive evolution in ecological time. Nat Ecol Evol. 2017 Oct; 1(10):1551–1561.

18. Frichot E, Francois O. LEA: an R package for Landscape and Ecological Association studies. Methods in Ecology and Evolution. 2015; http://membres-timc.imag.fr/Olivier.Francois/lea.html.

19. Gehan MA, Park S, Gilmour SJ, An C, Lee CM, Thomashow MF. Natural variation in the C-repeat binding factor cold response pathway correlates with local adaptation of Arabidopsis ecotypes. Plant J. 2015 Nov; 84(4):682–693.

20. Gloss AD, Vergnol A, Morton TC, Laurin PJ, Roux F, Bergelson J. Genome-wide association mapping within a local *Arabidopsis thaliana* population more fully reveals the genetic architecture for defensive metabolite diversity. Philos Trans R Soc Lond B Biol Sci. 2022 Jul; 377(1855):20200512.

21. Guo X, Liang R, Lou S, Hou J, Chen L, Liang X, Feng X, Yao Y, Liu J, Liu H. Natural variation in the SVP contributes to the pleiotropic adaption of *Arabidopsis thaliana* across contrasted habitats. J Genet Genomics. 2023 Dec; 50(12):993–1003.

22. Hancock AM, Brachi B, Faure N, Horton MW, Jarymowycz LB, Sperone FG, Toomajian C, Roux F, Bergelson J. Adaptation to climate across the *Arabidopsis thaliana* genome. Science. 2011; 334(6052):83–86.

23. Hoban S, Kelley JL, Lotterhos KE, Antolin MF, Bradburd G, Lowry DB, Poss ML, Reed LK, Storfer A, Whitlock MC. Finding the genomic basis of local adaptation: Pitfalls, practical solutions, and future directions. Am Nat. 2016 Oct; 188(4):379–397.

24. Horton MW, Hancock AM, Huang YS, Toomajian C, Atwell S, Auton A, Muliyati NW, Platt A, Sperone FG, Vilhjálmsson BJ, Nordborg M, Borevitz JO, Bergelson J. Genome-wide patterns of genetic variation in worldwide *Arabidopsis thaliana* accessions from the RegMap panel. Nat Genet. 2012; 44(2):212–216.

25. Huber CD, Nordborg M, Hermisson J, Hellmann I. Keeping It Local: Evidence for Positive Selection in Swedish *Arabidopsis thaliana*. Mol Biol Evol. 2014 Nov; 31(11):3026–3039.

26. Igolkina A, Bezlepsky A, Senti K, Nordborg M. Pannagram: unbiased pangenome alignment and mobilome calling across insects extends horizontal transfer from transposons to polintons. bioRxiv. 2026; p. 2025.02.07.637071.

27. Katz E, Li JJ, Jaegle B, Ashkenazy H, Abrahams SR, Bagaza C, Holden S, Pires CJ, Angelovici R, Kliebenstein DJ. Genetic variation, environment and demography intersect to shape Arabidopsis defense metabolite variation across Europe. Elife. 2021 May; 10.

28. Kerdaffrec E, Filiault DL, Korte A, Sasaki E, Nizhynska V, Seren Ü, Nordborg M. Multiple alleles at a single locus control seed dormancy in Swedish Arabidopsis. Elife. 2016 Dec; 5:e22502.

29. Kliebenstein DJ, Lambrix VM, Reichelt M, Gershenzon J, Mitchell-Olds. Gene duplication in the diversification of secondary metabolism: tandem 2-oxoglutarate-dependent dioxygenases control glucosinolate biosynthesis in Arabidopsis. Plant Cell. 2001; 13(3):681–693.

30. Lee G, Sanderson BJ, Ellis TJ, Dilkes BP, McKay JK, Ågren J, Oakley CG. A large-effect fitness trade-off across environments is explained by a single mutation affecting cold acclimation. Proc Natl Acad Sci U S A. 2024 Feb; 121(6):e2317461121.

31. Lee JH, Ryu HS, Chung KS, Posé D, Kim S, Schmid M, Ahn JH. Regulation of temperature-responsive flowering by MADS-box transcription factor repressors. Science. 2013; 342(6158):628–632.

32. Lo CY, Chien CC, Lin YP, Yeh PM, Lee CR. How weedy *Arabidopsis thaliana* dominated the world: ancestral variation and polygenic adaptation. bioRxiv. 2024 Apr; p. 2024.04.28.591542.

33. Long Q, Rabanal FA, Meng D, Huber CD, Farlow A, Platzer A, Zhang Q, Vilhjálmsson BJ, Korte A, Nizhynska V, Voronin V, Korte P, Sedman L, Mandáková T, Lysak MA, Seren U, Hellmann I, Nordborg M. Massive genomic variation and strong selection in *Arabidopsis thaliana* lines from Sweden. Nat Genet. 2013; 45(8):884–890.

34. de la Mata R, Mollá-Morales A, Méndez-Vigo B, Torres-Pérez R, Oliveros JC, Gómez R, Marcer A, Castilla AR, Nordborg M, Alonso-Blanco C, Picó FX. Variation and plasticity in life-history traits and fitness of wild *Arabidopsis thaliana* populations are not related to their genotypic and ecological diversity. BMC Ecology and Evolution. 2024 May; 24(1):1–18.

35. Méndez-Vigo B, Martínez-Zapater JM, Alonso-Blanco C. The Flowering Repressor SVP Underlies a Novel *Arabidopsis thaliana* QTL Interacting with the Genetic Background. PLoS Genet. 2013; 9(1):e1003289.

36. Nordborg M. Linkage disequilibrium, gene trees and selfing: an ancestral recombination graph with partial self-fertilization. Genetics. 2000 Feb; 154(2):923–929.

37. Nordborg M, Hu TT, Ishino Y, Jhaveri J, Toomajian C, Zheng H, Bakker E, Calabrese P, Gladstone J, Goyal R, Jakobsson M, Kim S, Morozov Y, Padhukasahasram B, Plagnol V, Rosenberg Na, Shah C, Wall JD, Wang J, Zhao K, et al. The pattern of polymorphism in *Arabidopsis thaliana*. PLoS Biol. 2005 Jul; 3(7):e196.

38. Oakley CG, Schemske DW, McKay JK, Ågren J. Ecological genetics of local adaptation in *Arabidopsis*: An 8-year field experiment. Mol Ecol. 2023 Jun; .

39. Ohta T. Linkage disequilibrium due to random genetic drift in finite subdivided populations. Proc Natl Acad Sci U S A. 1982 Mar; 79(6):1940–1944.

40. Pagnussat GC, Yu HJ, Ngo QA, Rajani S, Mayalagu S, Johnson CS, Capron A, Xie LF, Ye D, Sundaresan V. Genetic and molecular identification of genes required for female gametophyte development and function in Arabidopsis. Development. 2005 Feb; 132(3):603–614.

41. Paradis E, Schliep K. ape 5.0: an environment for modern phylogenetics and evolutionary analyses in R. Bioinformatics. 2019; 35:526–528. doi: 10.1093/bioinformatics/bty633.

42. Pau G, Fuchs F, Sklyar O, Boutros M, Huber W. EBImage–an R package for image processing with applications to cellular phenotypes. Bioinformatics. 2010 Apr; 26(7):979–981.

43. Pisupati R, Reichardt I, Seren Ü, Korte P, Nizhynska V, Kerdaffrec E, Uzunova K, Rabanal FA, Filiault DL, Nordborg M. Verification of Arabidopsis stock collections using SNPmatch, a tool for genotyping high-plexed samples. Sci Data. 2017 Dec; 4:170184.

44. Platt A, Horton M, Huang YS, Li Y, Anastasio AE, Mulyati NW, Agren J, Bossdorf O, Byers D, Donohue K, Dunning M, Holub EB, Hudson A, Le Corre V, Loudet O, Roux F, Warthmann N, Weigel D, Rivero L, Scholl R, et al. The scale of population structure in *Arabidopsis thaliana*. PLoS Genet. 2010; 6(2):e1000843.

45. Platt A, Vilhjálmsson BJ, Nordborg M. Conditions under which genome-wide association studies will be positively misleading. Genetics. 2010 Nov; 186(3):1045–1052.

46. Postma FM, Ågren J. Early life stages contribute strongly to local adaptation in *Arabidopsis thaliana*. Proc Natl Acad Sci U S A. 2016 Jul; 113(27):7590–7595.

47. Rockman MV. The QTN program and the alleles that matter for evolution: all that’s gold does not glitter. Evolution. 2012 Jan; 66(1):1–17.

48. Savolainen O, Lascoux M, Merilä J. Ecological genomics of local adaptation. Nat Rev Genet. 2013 Nov; 14(11):807–820.

49. Schmitz G, Linstädter A, Frank ASK, Dittberner H, Thome J, Schrader A, Linne von Berg KH, Fulgione A, Coupland G, de Meaux J. Environmental filtering of life-history trait diversity in urban populations of *Arabidopsis thaliana*. J Ecol. 2024 Jan; 112(1):14–27.

50. Slovak R, Setzer C, Roiuk M, Bertels J, Göschl C, Jandrasits K, Beemster GTS, Busch W. Ribosome assembly factor Adenylate Kinase 6 maintains cell proliferation and cell size homeostasis during root growth. New Phytol. 2020 Mar; 225(5):2064–2076.

51. Sork VL. Genomic studies of local adaptation in natural plant populations. J Hered. 2017 Dec; 109(1):3–15.

52. Stratton DA. Genotype-by-environment interactions for fitness of *Erigeron annuus* show fine-scale selective heterogeneity. Evolution. 1994 Oct; 48(5):1607–1618.

53. Stratton DA. Spatial Scale of Variation in Fitness of *Erigeron annuus*. Am Nat. 1995; 146(4):608–624.

54. The 1001G+ Consortium. The 1001G+ project: A curated collection of *Arabidopsis thaliana* long-read genome assemblies to advance plant research. bioRxiv. 2024 Dec; p. 2024.12.23.629943.

55. Travisano M, Shaw RG. Lost in the map. Evolution. 2013 Feb; 67(2):305–314.

56. Turesson G. The genotypical response of the plant species to the habitat. Hereditas. 1922; 3:211–350.

57. Vilhjálmsson BJ, Nordborg M. The nature of confounding in genome-wide association studies. Nat Rev Genet. 2012; 14(1):1–2.

58. Wahlund S. Zusammensetzung von Populationen und Korrelationserscheinungen vom Standpunkt der Vererbungslehre aus betrachtet. Hereditas. 1928; 11(1):65–106.

59. Yin L, Zhang H, Tang Z, Xu J, Yin D, Zhang Z, Yuan X, Zhu M, Zhao S, Li X, Liu X. rMVP: A Memory-efficient, Visualization-enhanced, and Parallel-accelerated Tool for Genome-wide Association Study. Genomics Proteomics Bioinformatics. 2021 Aug; 19(4):619–628.

60. Yu J, Pressoir G, Briggs WH, Vroh Bi I, Yamasaki M, Doebley JF, McMullen MD, Gaut BS, Nielsen DM, Holland JB, Kresovich S, Buckler ES. A unified mixed-model method for association mapping that accounts for multiple levels of relatedness. Nat Genet. 2006 Feb; 38(2):203–208.

61. Zhao K, Aranzana MJ, Kim S, Lister C, Shindo C, Tang C, Toomajian C, Zheng H, Dean C, Marjoram P, Nordborg M. An Arabidopsis example of association mapping in structured samples. PLoS Genet. 2007; 3(1):e4.

62. Zhao Y, Christensen SK, Fankhauser C, Cashman JR, Cohen JD, Weigel D, Chory J. A role for flavin monooxygenase-like enzymes in auxin biosynthesis. Science. 2001 Jan; 291(5502):306–309.

63. Zou YP, Hou XH, Wu Q, Chen JF, Li ZW, Han TS, Niu XM, Yang L, Xu YC, Zhang J, Zhang FM, Tan D, Tian Z, Gu H, Guo YL. Adaptation of *Arabidopsis thaliana* to the Yangtze River basin. Genome Biol. 2017 Dec; 18(1):239.

