## Supplementary material for "Life-history trade-offs explain local adaptation in *Arabidopsis thaliana*": Figure supplements

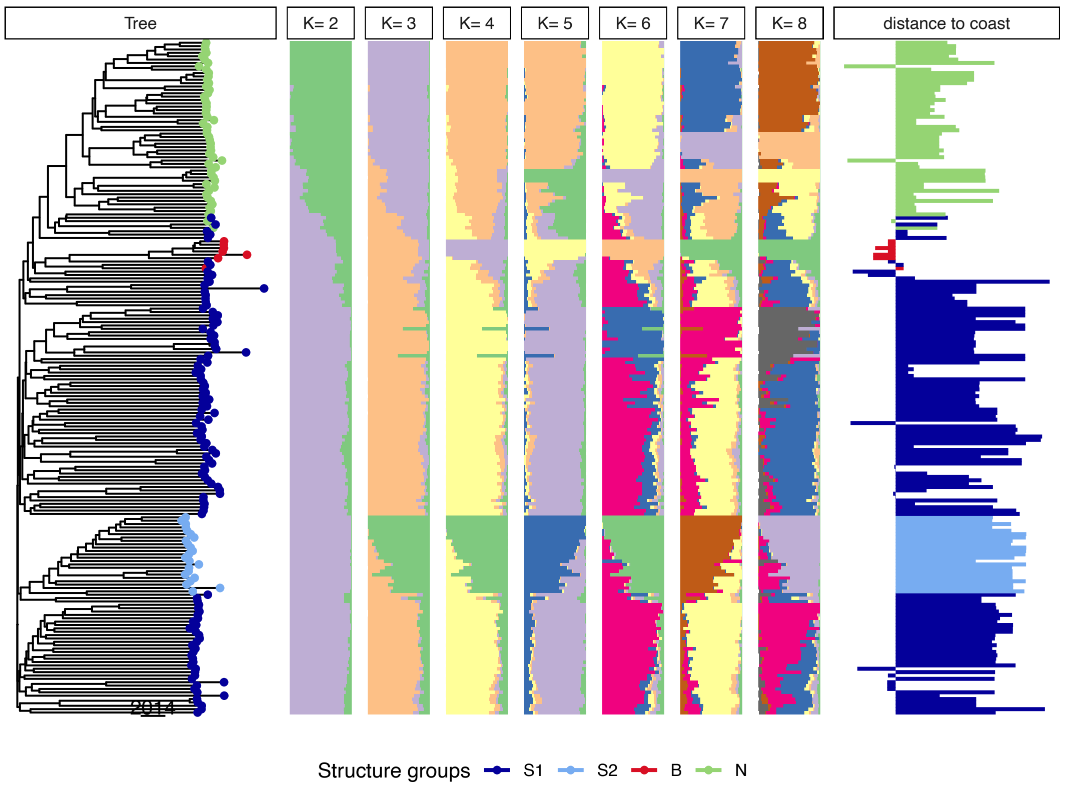


**Figure 1—figure supplement 1.** Population structure in the 200 Swedish accessions. On the left is a Neighbor-Joining tree of all accessions included in this study. Tree tips are colored according to the structure groups used in this paper. The genetic distance between accessions was computed using 124,071 LD-pruned SNPs. The following panels show ADMIXTURE results for different values of $K$. Note that for $K > 4$, only “admixed" colors are added. The right-most panel shows the distance to the Baltic Sea for each accession, in hundreds of meters and presented on a $\log_{10}$ scale. Hence, leftward pointing bars highlight accessions collected less than 100 m from the coastline.

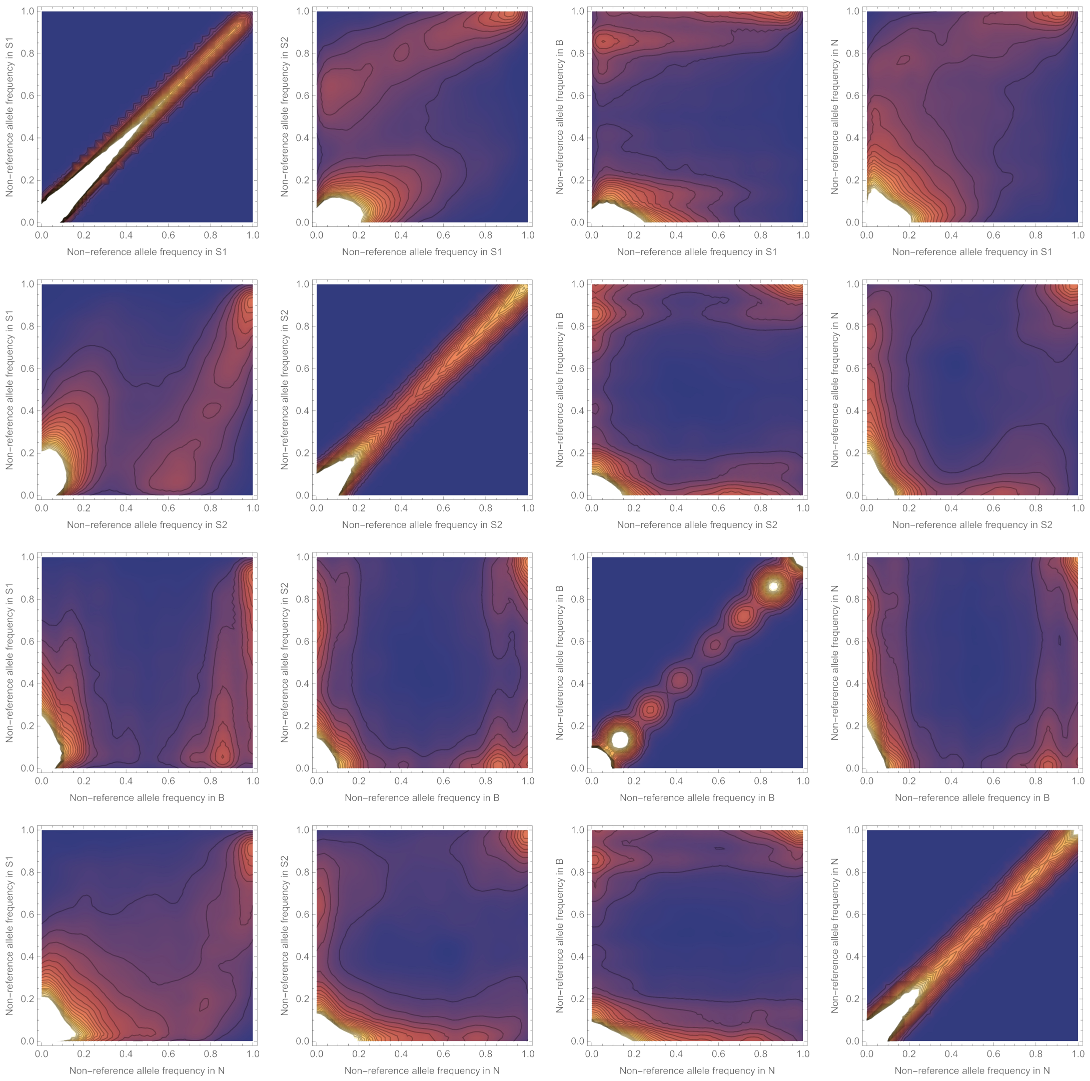


**Figure 1—figure supplement 2.** Density plot of the joint distribution of the non-reference SNP allele frequencies between the genetic groups. As expected from basic population genetics theory, the non-reference allele tends to be rare in all populations, and the plots have therefore been clipped (white color) to prevent the high density of SNPs close to the origin from obscuring more interesting parts of the distribution.

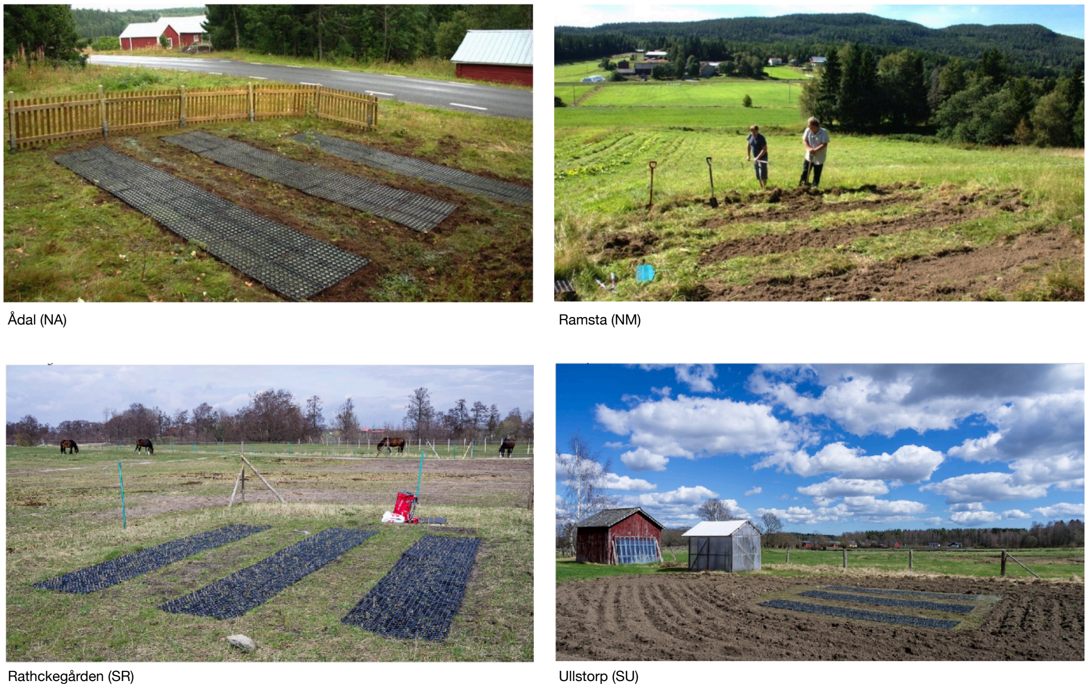


**Figure 1—figure supplement 3.** Photos of the four common-garden field sites: SR and SU in the south; NA and NM in the north. People in NM photo are authors M.B. and S.H. Each experiment consisted consisted three blocks of eight replicates of the 200 accessions, laid across the most obvious environmental gradient (distance to road, slope, distance from trees, *etc*).

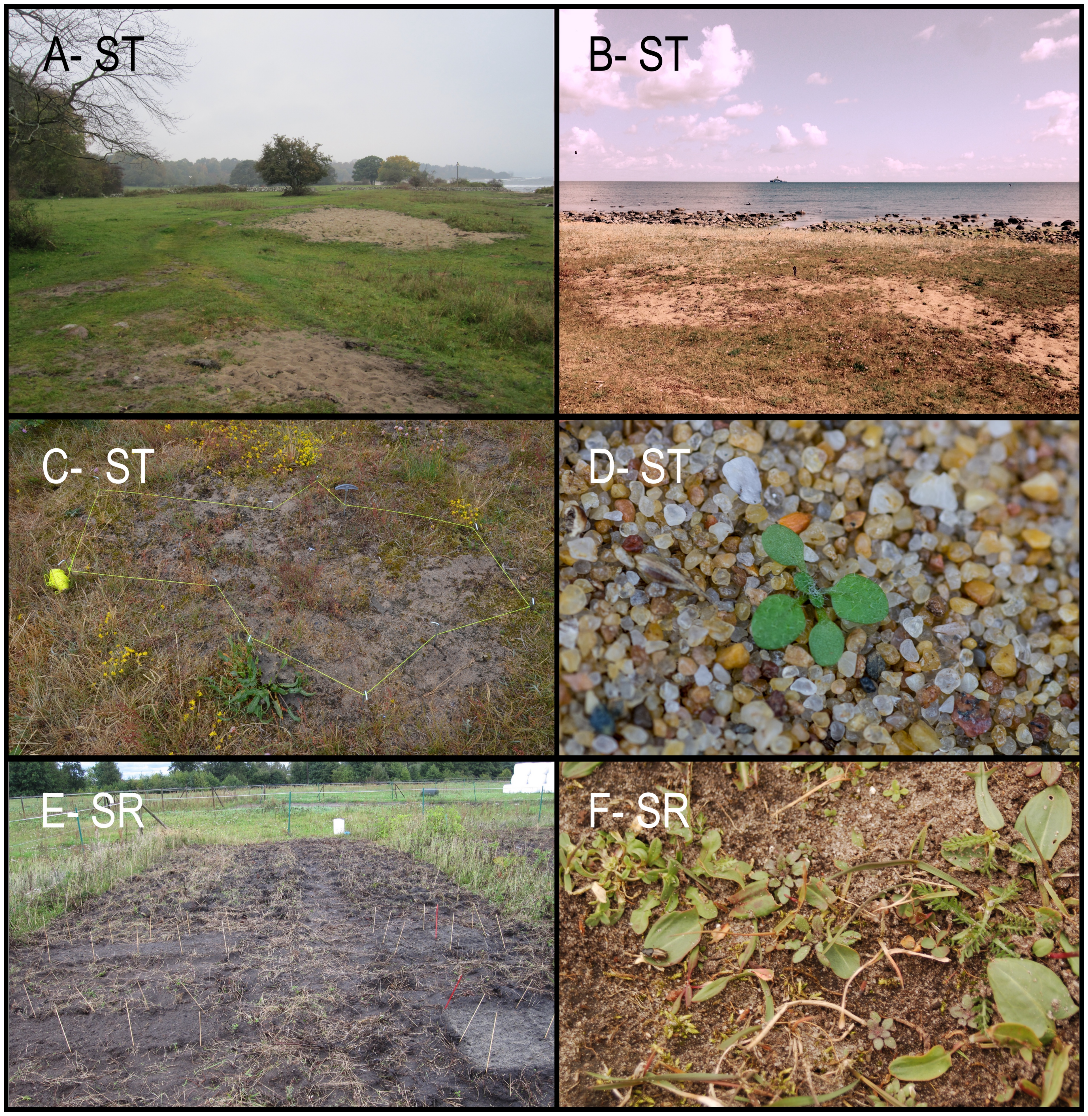


**Figure 1—figure supplement 4.** Selection experiment field sites in southern Sweden and sampling strategy. (**A**) Overview of the ST site, a cow pasture on a beach by the Baltic Sea. (**B**) Another view of ST showing its proximity to the sea. (**C**) An experimental plot in ST, outlined with yellow string. Each plot was originally ≈1 m$^2$, comprised of four 60 × 40 cm subplots placed to include suitable open habitat. The subplots were marked with metal nails pushed into the ground. When sampling, we first used a metal detector to find the nails and relocated the subplots, then searched the surrounding area for plants to assess how far they had spread since the original sowing, defining the new sampling area as illustrated in the photo. (**D**) An *A. thaliana* seedling growing in the sand at ST. (**E**) The SR site, an agricultural field that was tilled and weeded before the experiment was installed; sticks mark the arrangement of the 60 × 40 cm subplots. (**F**) *A. thaliana* growing at SR.

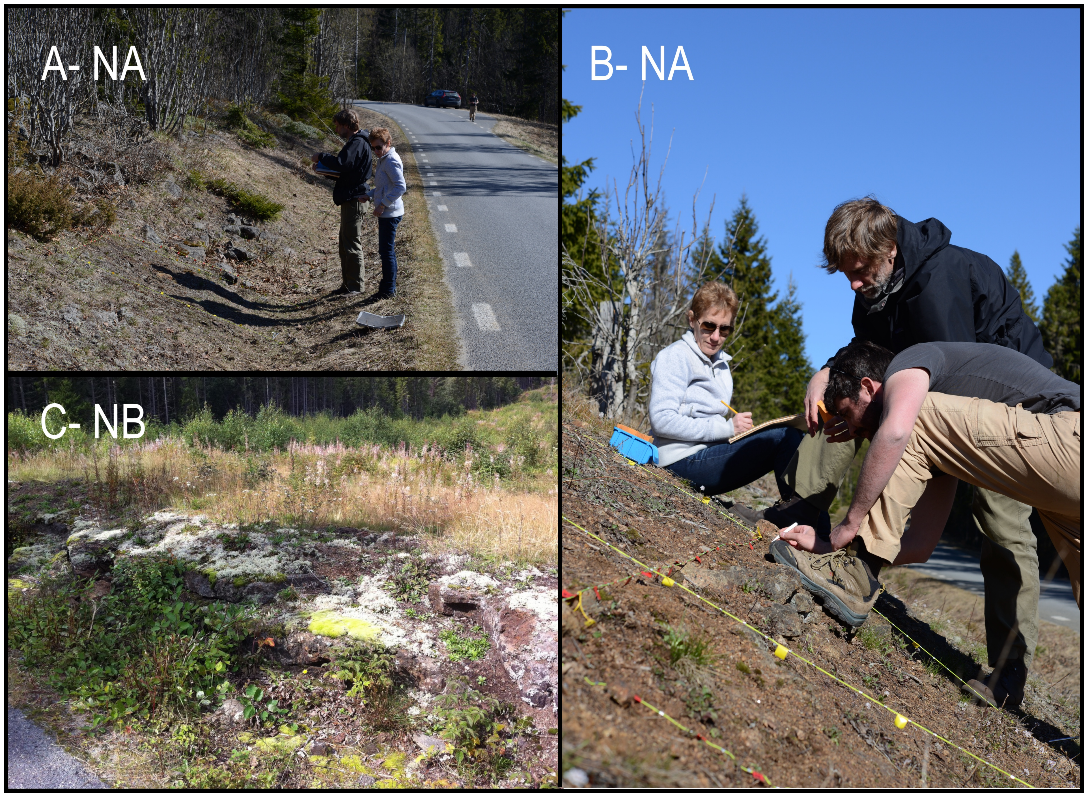


**Figure 1—figure supplement 5.** Selection experiment field sites in northern Sweden and sampling strategy. (**A**) The NA site, a south-facing, rocky, road-side slope being assessed by M.N. and J.B. (**B**) The NA site being sampled by J.B., B.B. and M.N. The plots were established using 60 × 40 cm subplots as described above, but as they expanded through seed dispersal, we developed a system to ensure random, unbiased sampling despite the irregular shape. After assessing the extent of the area occupied by plants around the initial plots, we installed two parallel strings on either side of the plot to create a coordinate system, visible here, within which transect positions could be randomly assigned. Along each transect, plants were sampled at regular 5 cm intervals, always selecting the plant whose center fell closest to the mark. Sampling continued until reaching our target of 70 individuals. (**C**) The NB site, a very similar, south-facing, rocky slope near a road.

980

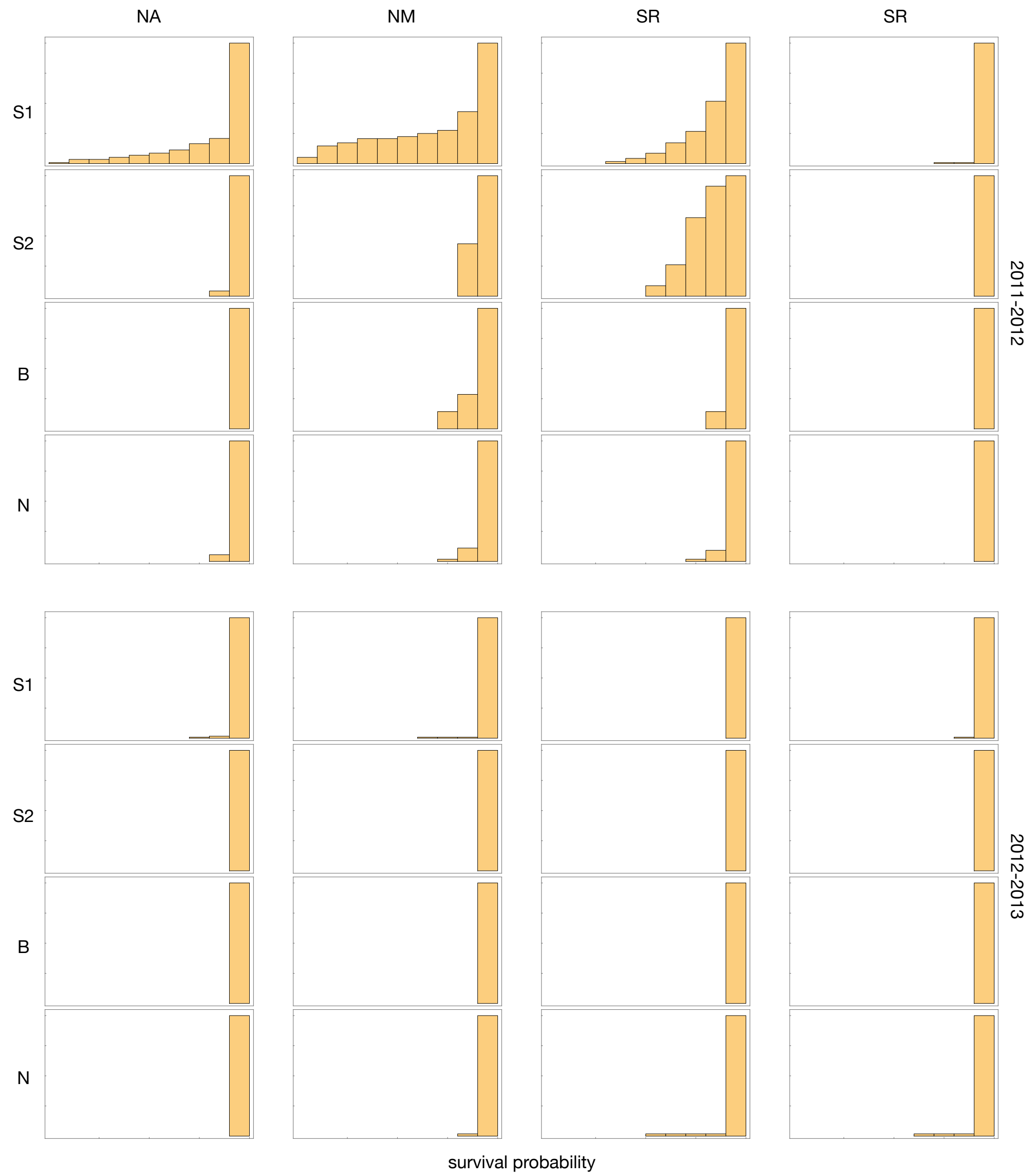


**Figure 2—figure supplement 1.** Histograms showing the cumulative density of survival estimates across sites and years and groups. In 2011-12, the two northern sites saw high mortality primarily of S1 accessions, and one of the southern sites, SR, saw high mortality primarily of S1 and S2 accession. Mortality in the remaining five experiments was trivial.

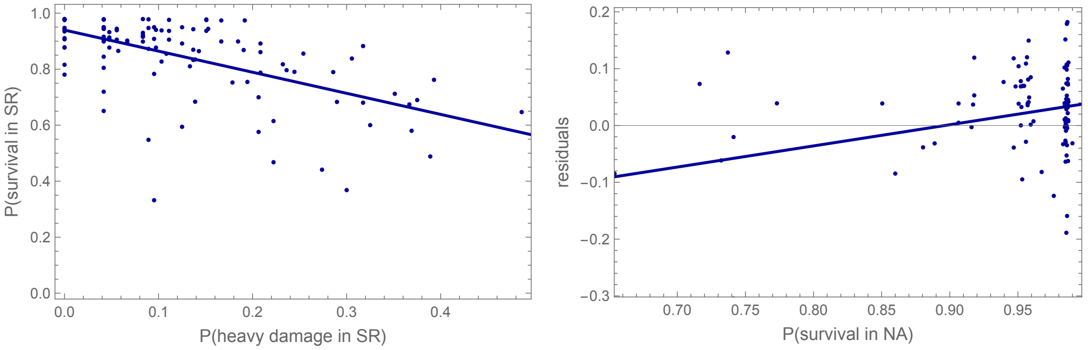


**Figure 4—figure supplement 1.** Left: Overwinter survival in SR 2011-12 was predicted by slug damage ($R^2_{\text{adj.}} = 0.33; p = 8.6 \times 10^{-12}$). Right: residuals from regression of survival in SR on slug damage remain positively correlated with survival in NA 2011-12, suggesting an underlying shared cause ($R^2_{\text{adj.}} = 0.37; p = 9.3 \times 10^{-14}$).

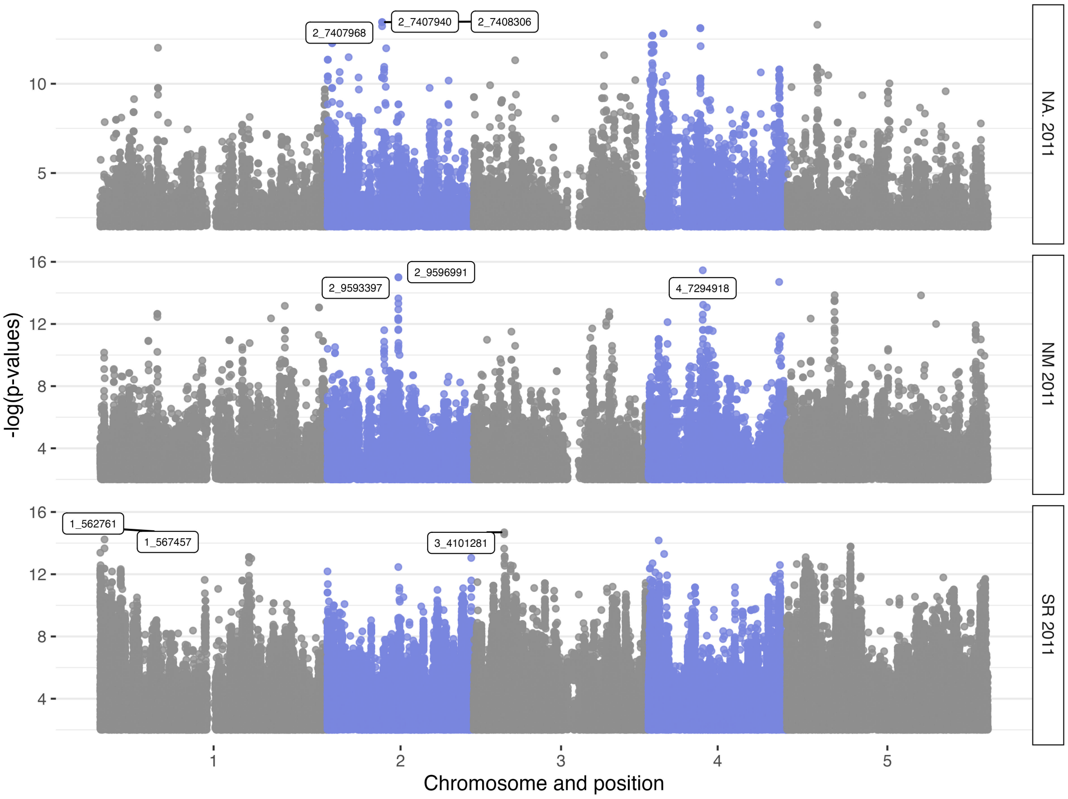


**Figure 5—figure supplement 1.** Manhattan plots of GWAS for overwinter survival without structure correction.

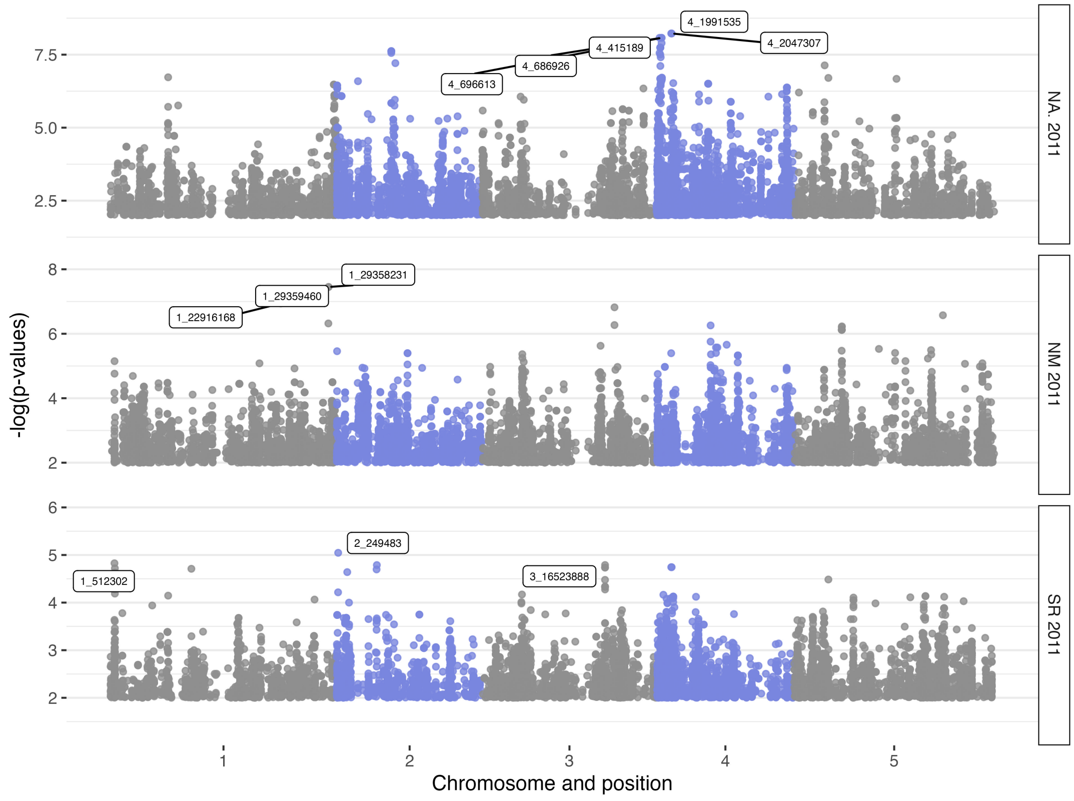


**Figure 5—figure supplement 2.** Manhattan plots of GWAS for overwinter survival with structure correction.

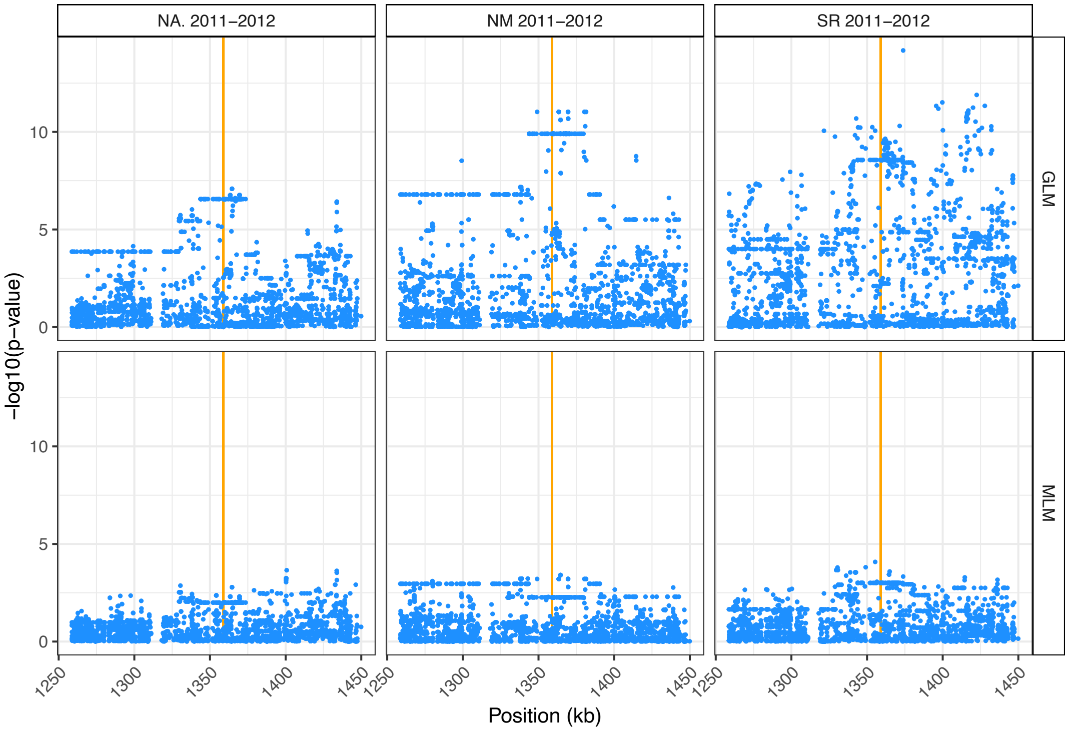


**Figure 5—figure supplement 3.** Zoom-in on *AOP* region on chromosome 4, without (top row; *cf*. ***Figure 5—figure Supplement 1***) and with (top row; *cf*. ***Figure 5—figure Supplement 2***) structure correction. The orange line shows the location of the gene.

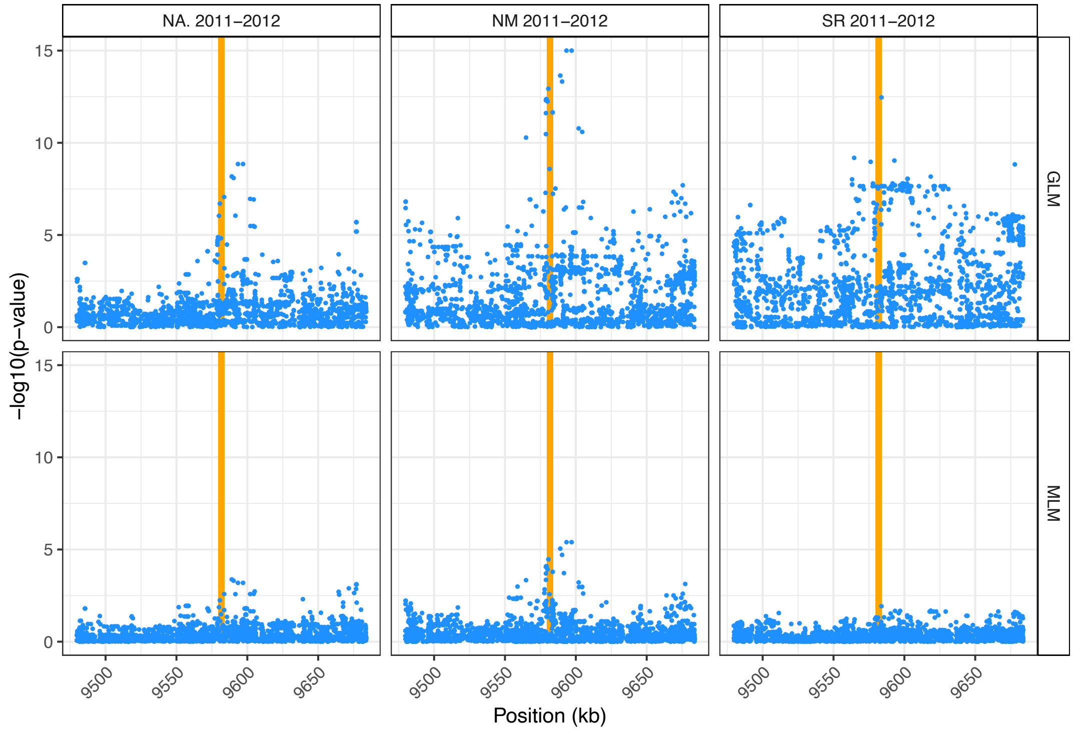


**Figure 5—figure supplement 4.** Zoom-in on *SVP* region on chromosome 2, without (top row; *cf*. ***Figure 5—figure Supplement 1***) and with (bottom row; *cf*. ***Figure 5—figure Supplement 2***) structure correction. The orange line shows the location of the gene.

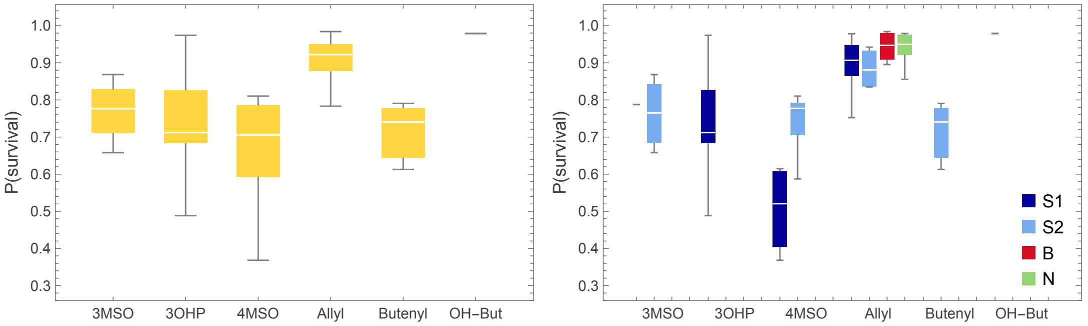


**Figure 6—figure supplement 1.** Survival was also associated with glucosinolate profiles. Same plots as in ***Figure 6***, but with over-winter survival as dependent variable.

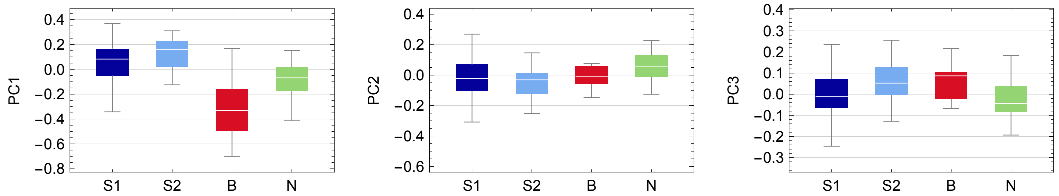


**Figure 8—figure supplement 1.** Box plots showing the distribution of each PC across accession by group.

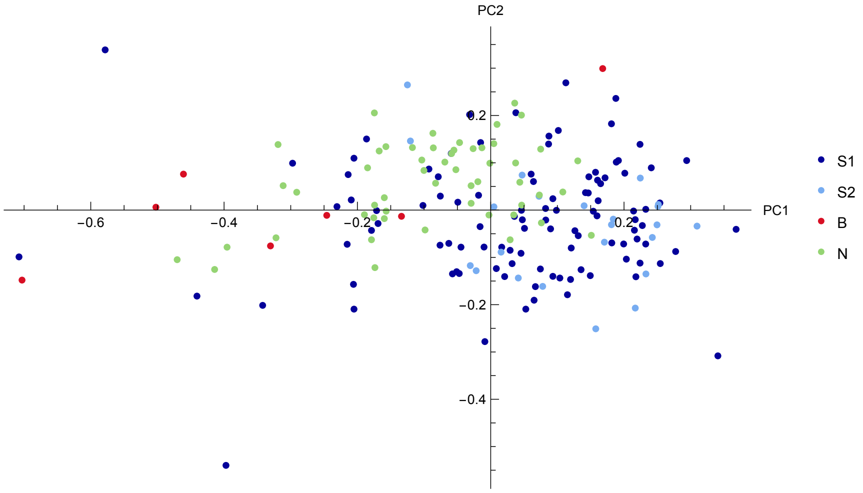


**Figure 8—figure supplement 2.** Scatter plot showings the accessions, colored by group, in the PC1-PC2 coordinate system.

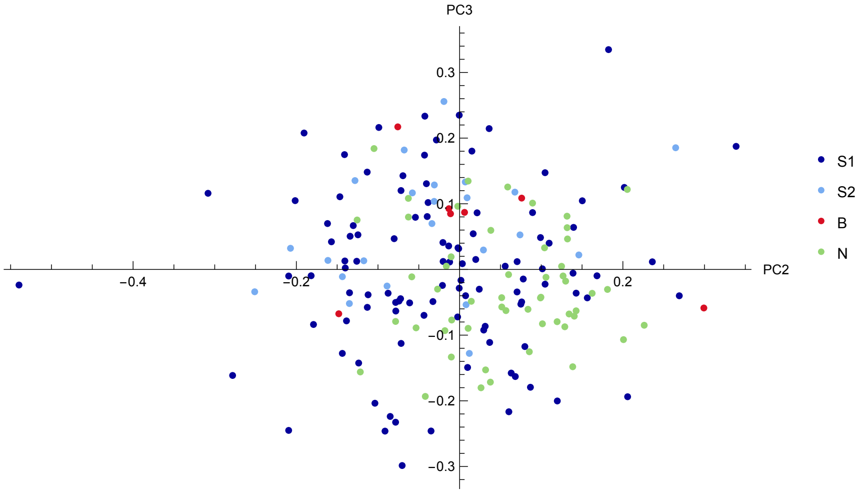


**Figure 8—figure supplement 3.** Scatter plot showings the accessions, colored by group, in the PC2-PC3 coordinate system.

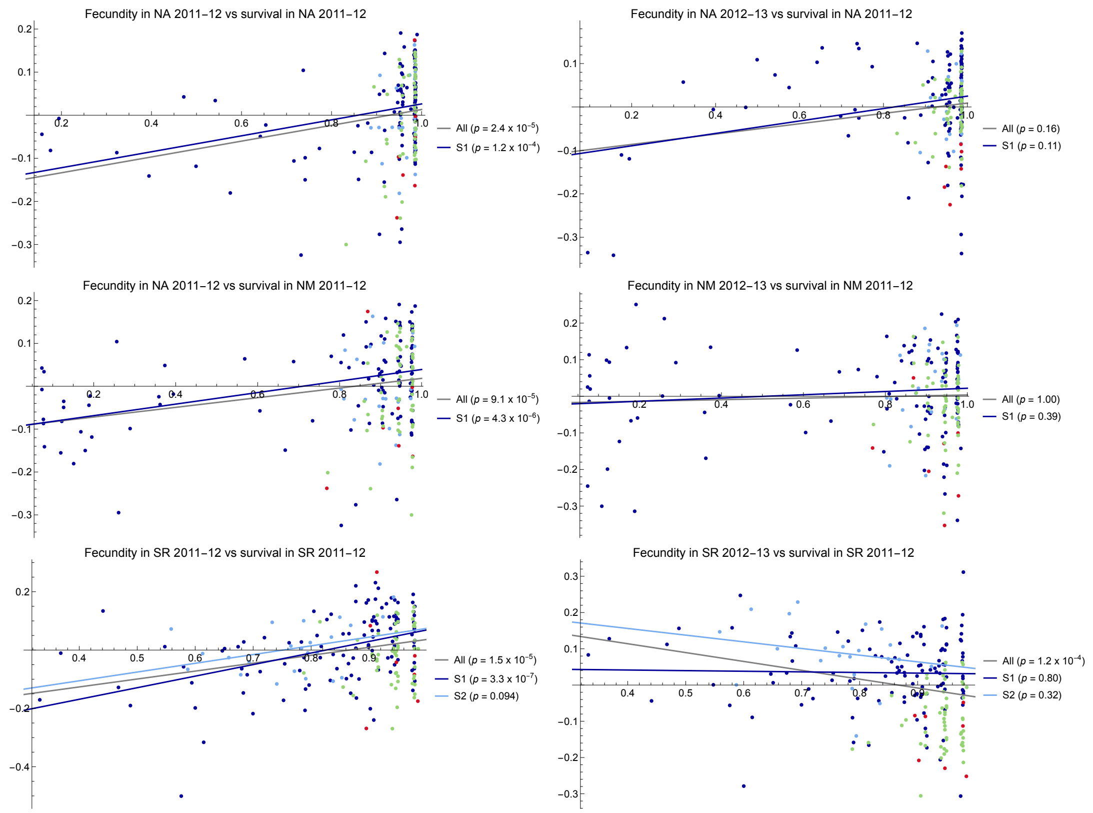


**Figure 9—figure supplement 1.** The left column shown that overwinter survival and springtime fecundity among survivors is significantly positively correlated in all three experiments with non-trivial mortality, suggesting that a common factor is influencing both. The right column uses fecundity in the following season as a control to demonstrate that no correlation (within groups) exist when no direct causation is possible.

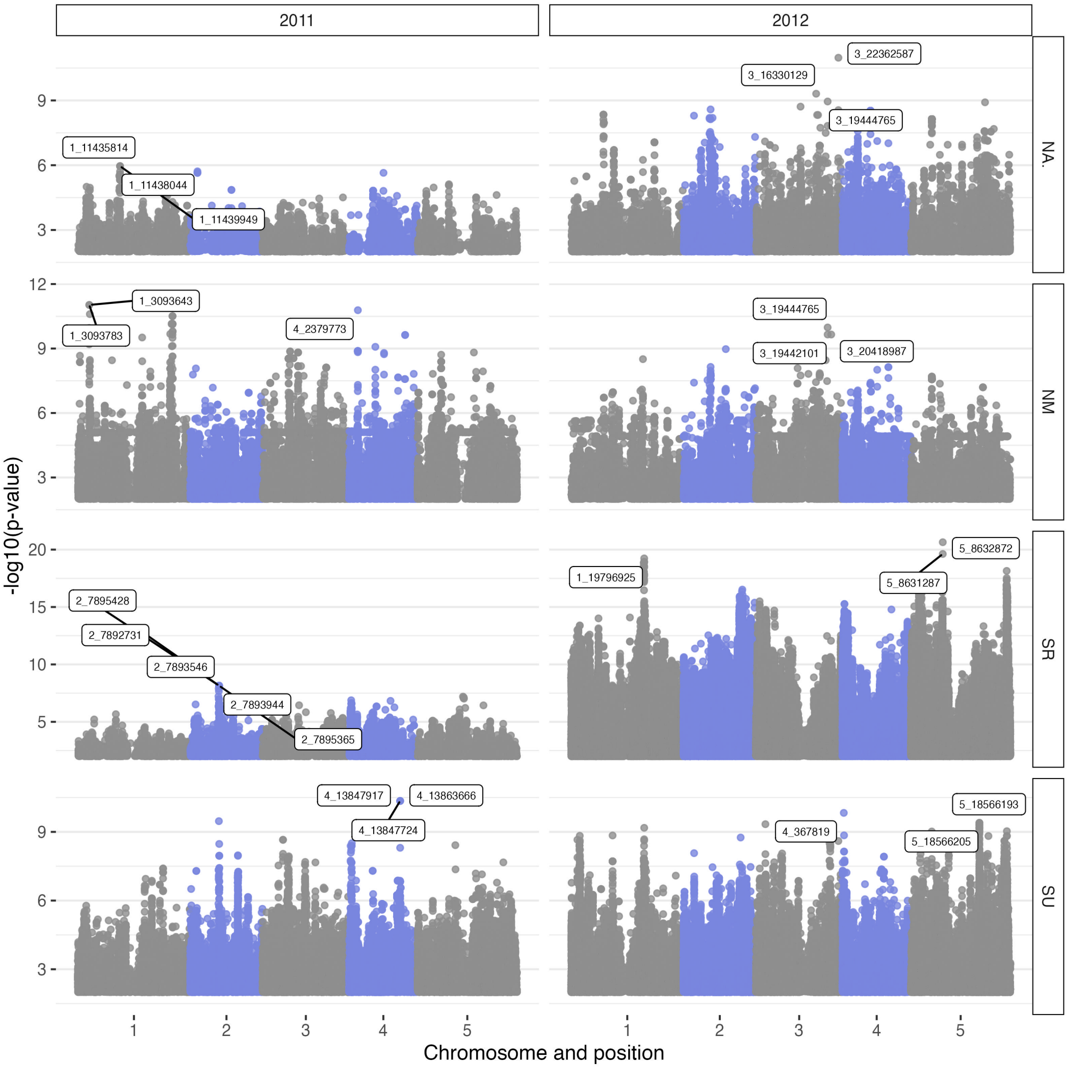


**Figure 12—figure supplement 1.** Manhattan plots for fecundity GWAS without correction for structure.

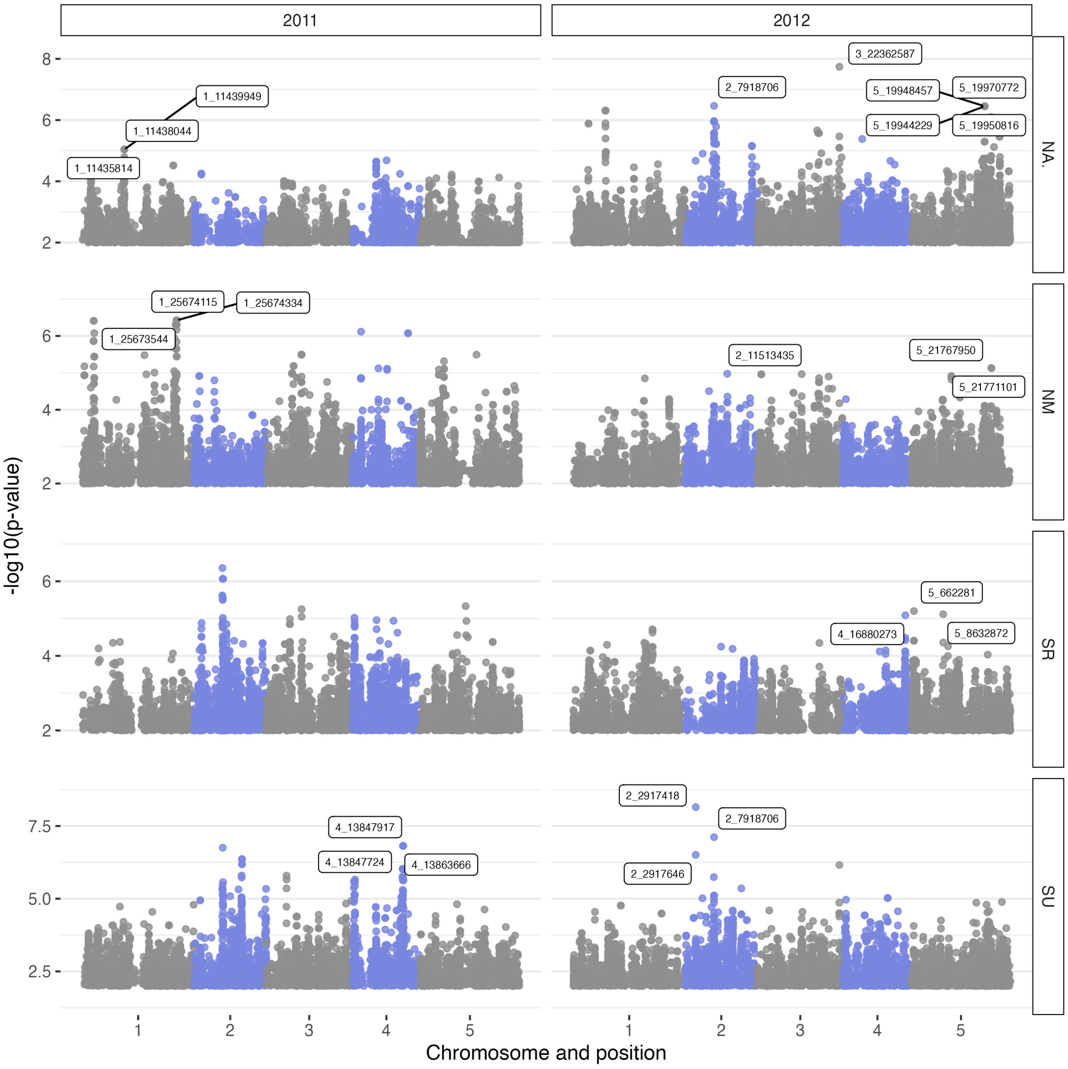


**Figure 12—figure supplement 2.** Manhattan plots for fecundity GWAS with correction for structure.

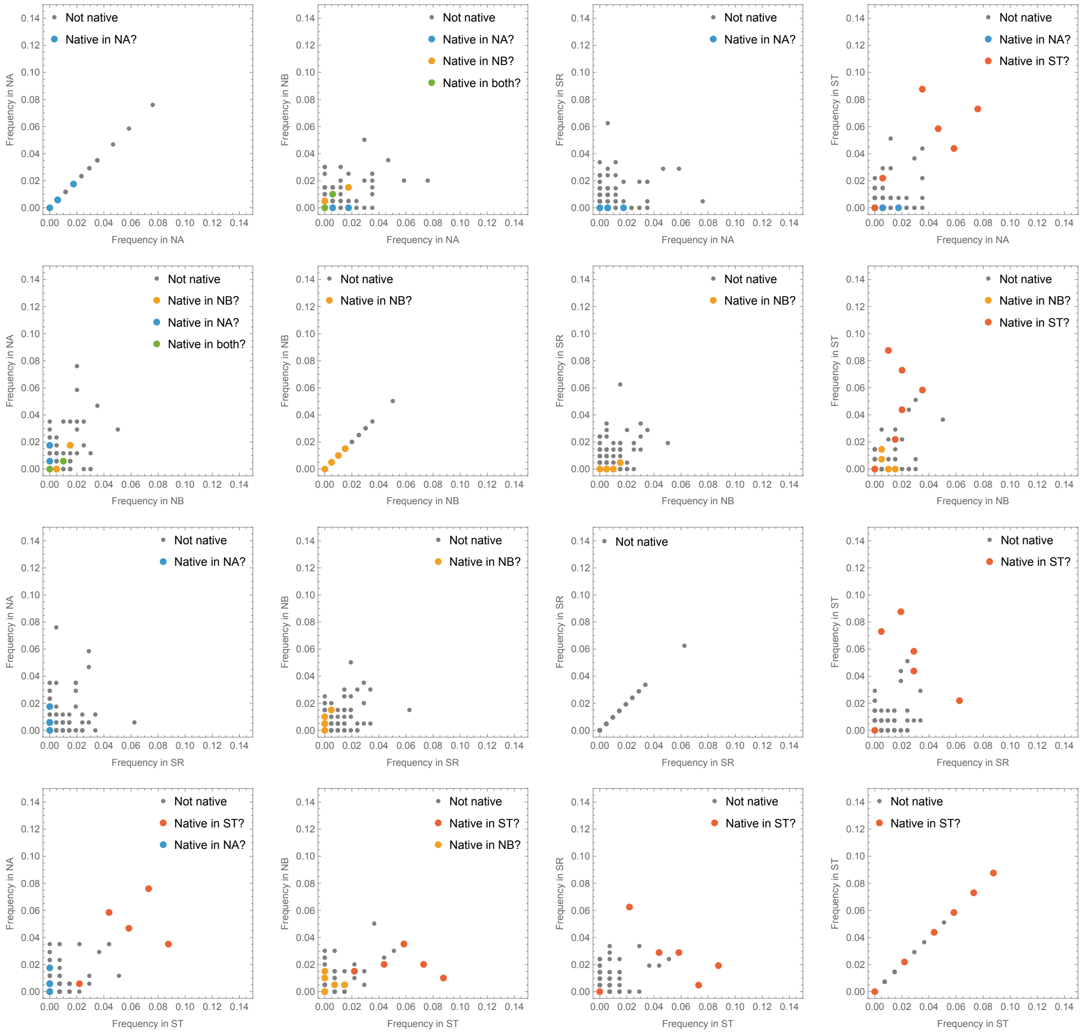


**Figure 13—figure supplement 1.** Same as ***Figure 13***, but with the two obvious outliers removed to show remaining data better.

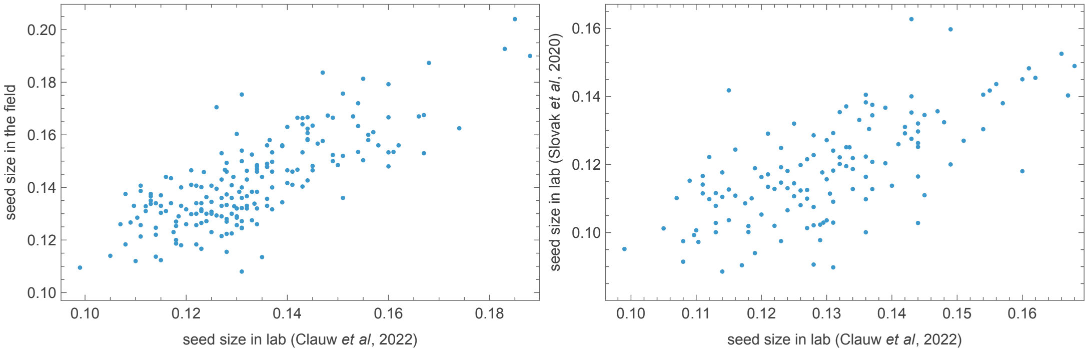


**Figure 15—figure supplement 1.** Correlation of seed size across environments. Lab measurements from ***Clauw et al.*** (***2022***) against unpublished field measurements, and against lab measurements from ***Slovak et al.*** (***2020***), respectively.

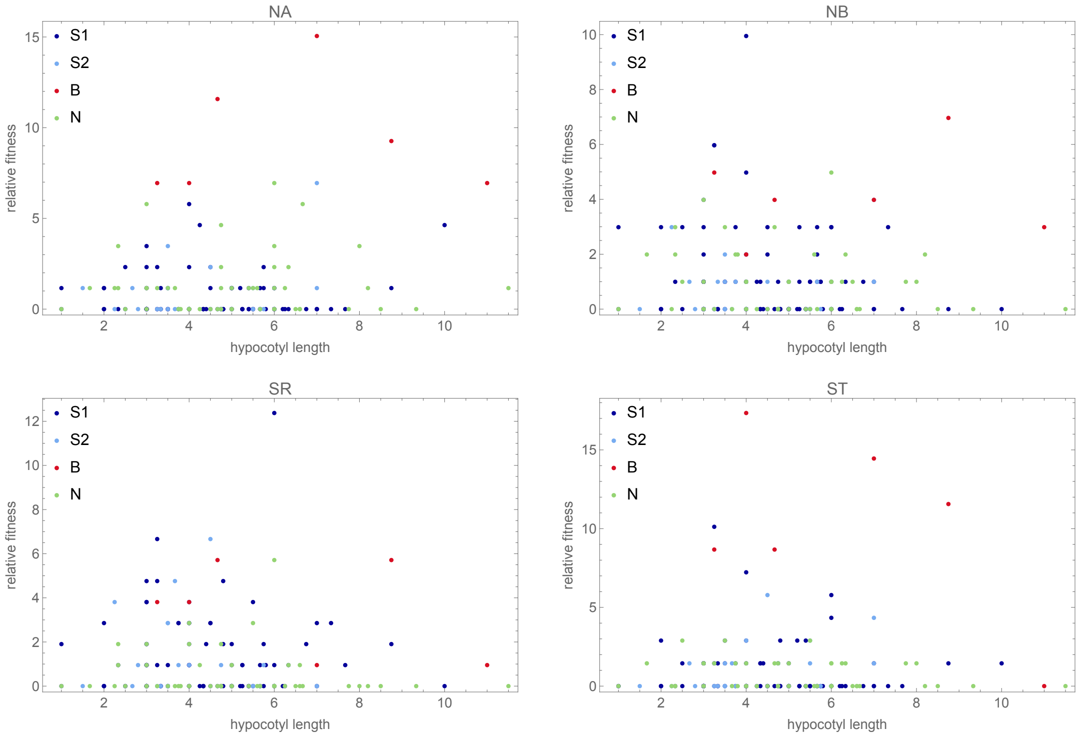


**Figure 16—figure supplement 1.** Fitness as a function of hypocotyl elongation in each experiment separately.

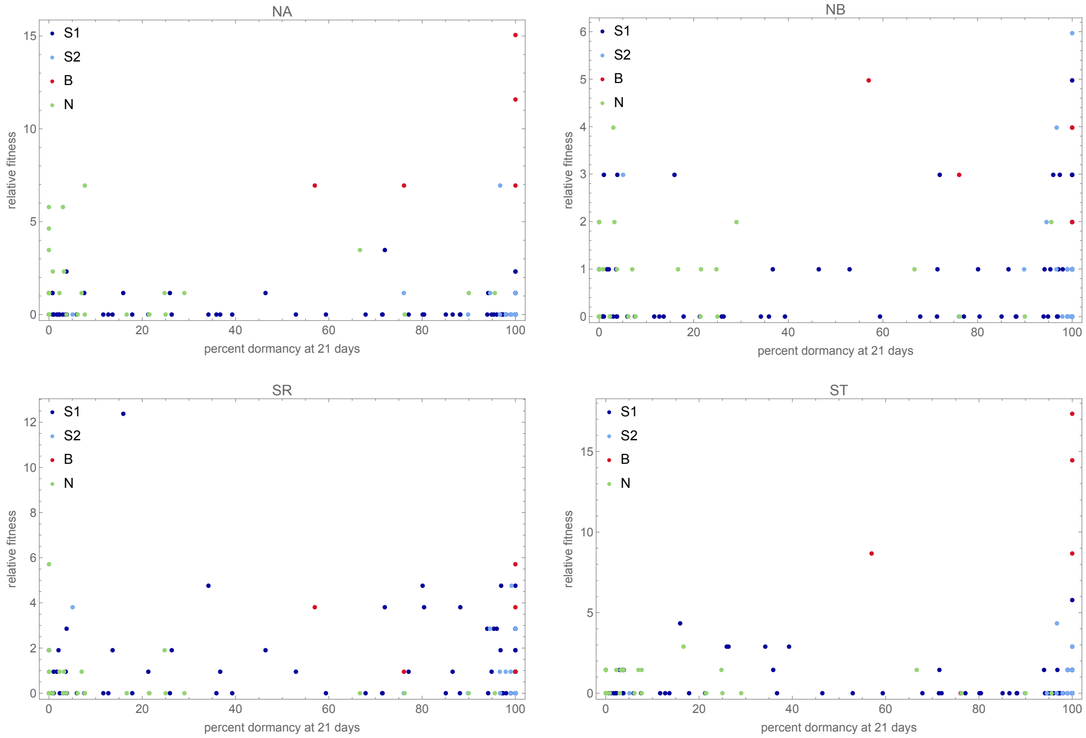


**Figure 16—figure supplement 2.** Fitness as a function of primary seed dormancy in each experiment separately. High dormancy is necessary (but not sufficient) for high fitness in the southern beach site (ST) but appears to be less important in the other sites, consistent with local adaptation (***Kerdaffrec et al., 2016***).

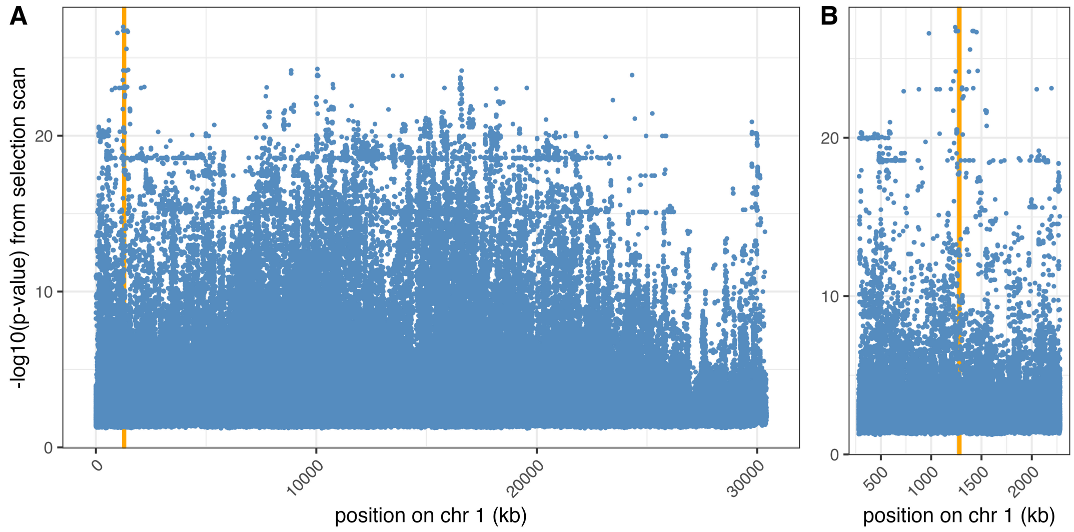


**Figure 18—figure supplement 1.** Selection scan peak on chromosome 1. Orange vertical lines indicates the position of *YUC3* (AT1G04610) and *MEE4* (AT1G04630). (**A**) Whole chromosome. (**B**) Zoom-in on peak.

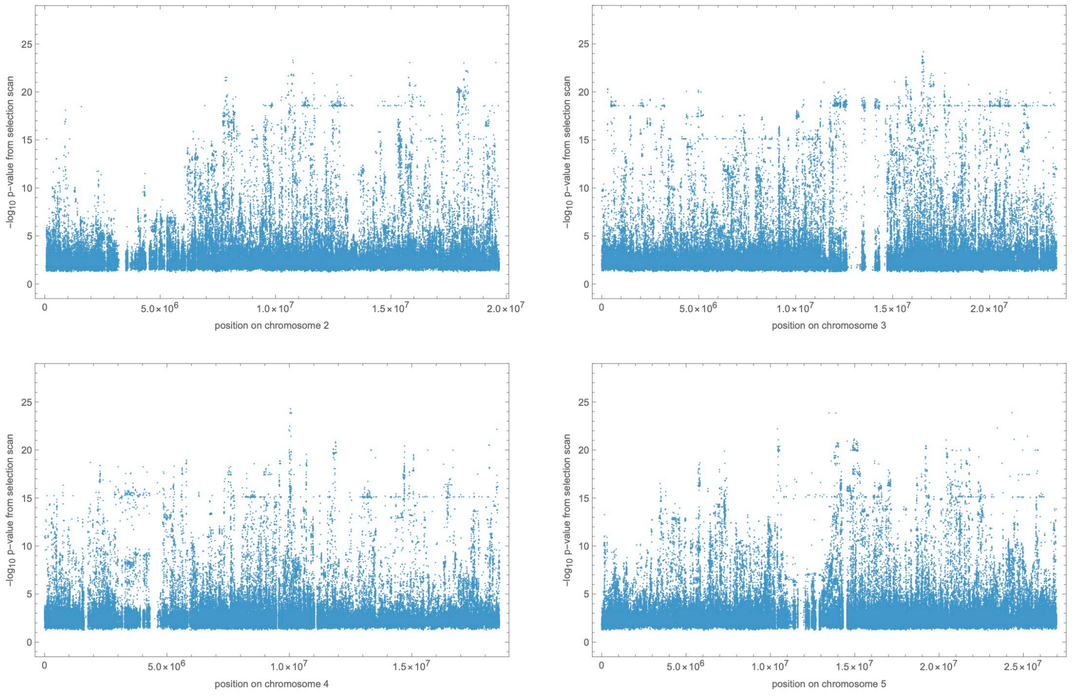


**Figure 18—figure supplement 2.** Result of selection scan for chromosomes 2–4. Negative log p-values have been summed across the four experiments. For chromosome 1, see ***Figure 18***.

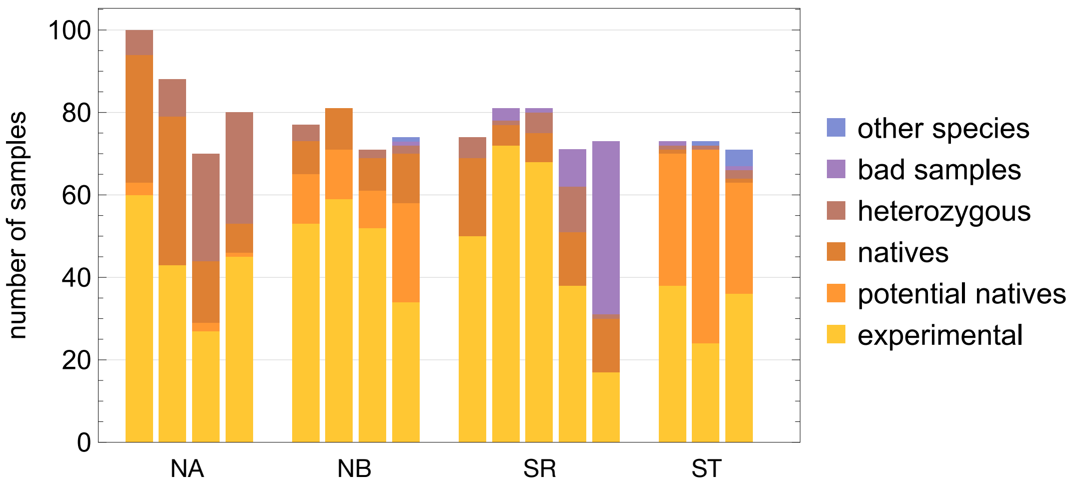


**Figure 20—figure supplement 1.** The number of samples in each category per site and plot. A minimum of 70 plants were sample per surviving plot.
